# A sort-seq approach to the development of single fluorescent protein biosensors

**DOI:** 10.1101/2020.08.21.261578

**Authors:** John N. Koberstein, Melissa L. Stewart, Taylor L. Mighell, Chadwick B. Smith, Michael S. Cohen

## Abstract

The utility of single fluorescent protein biosensors (SFPBs) in biological research is offset by the difficulty in engineering these tools. SFPBs generally consist of three basic components: a circularly permuted fluorescent protein, a ligand-binding domain, and a pair of linkers connecting the two domains. In the absence of predictive methods for biosensor engineering, most designs combining these three components will fail to produce allosteric coupling between ligand binding and fluorescence emission. Methods to construct libraries of biosensor designs with variations in the site of GFP insertion and linker sequences have been developed, however, our ability to construct new variants has exceeded our ability to test them for function. Here, we address this challenge by applying a massively parallel assay termed “sort-seq” to the characterization of biosensor libraries. Sort-seq combines binned fluorescence-activated cell sorting, next-generation sequencing, and maximum likelihood estimation to quantify the dynamic range of many biosensor variants in parallel. We applied this method to two common biosensor optimization tasks: choice of insertion site and optimization of linker sequences. The sort-seq assay applied to a maltose-binding protein domain-insertion library not only identified previously described high-dynamic-range variants but also discovered new functional insertion-sites with diverse properties. A sort-seq assay performed on a pyruvate biosensor linker library expressed in mammalian cell culture identified linker variants with substantially improved dynamic range. Machine learning models trained on the resulting data can predict dynamic range from linker sequence. This high-throughput approach will accelerate the design and optimization of SFPBs, expanding the biosensor toolbox.

## Introduction

Genetically-encoded single fluorescent protein biosensors (SFPBs) can unmask aspects of cellular signaling and metabolism that cannot be detected using traditional biochemical approaches (Okumoto et al., 2012). Biosensors can provide crucial information on the subcellular compartmentalization of analytes, resolve changes in concentration over time, and highlight cellular heterogeneity (Arce-Molina et al., 2020; Hou et al., 2011; Miyawaki and Niino, 2015; Ni et al., 2018; Okumoto et al., 2012; Pendin et al., 2017). However, each SFPB specifically measures a single analyte, requiring a unique biosensor for each target. A better understanding of the factors underlying successful designs is necessary to accelerate the development of novel biosensors.

SFPBs can be created by inserting a fluorescent protein (FP) into a ligand-binding domain (LBD) such that ligand binding allosterically regulates fluorescence (Arce-Molina et al., 2020; Doi and Yanagawa, 1999; Marvin et al., 2013, 2011; Nadler et al., 2016). The nature of this allosteric domain coupling is not well understood and has proven difficult to design rationally (Nadler et al., 2016). In the absence of a predictable relationship between biosensor sequence and function, low-throughput protein-engineering methods are unlikely to discover the optimal sequence because they can explore only a small subset of the possible design space. While the rules for combining domains to produce allostery in biosensors are unclear, the site of insertion and the amino acids connecting the domains are thought to be the two most important parameters (Marvin et al., 2011). As an alternative to rational design, unbiased library methods have been developed to enable the creation of many domain-insertion and linker sequence variants in parallel (Nadler et al., 2016). However, the major challenge is efficiently screening these variants.

One possible solution is to apply massively parallel assays that link cellular phenotype with DNA sequencing to characterize sequence-function pairs in a high-throughput fashion. A type of massively parallel assay combining fluorescence-activated cell sorting (FACS) with sequencing, referred to as sort-seq, is particularly relevant for screening biosensor libraries (Kinney et al., 2010; Kosuri et al., 2013; Noderer et al., 2014; Peterman et al., 2014; Peterman and Levine, 2016; Sharon et al., 2014, 2012). In the simplest sort-seq experimental design, the library is sorted to collect cells above a threshold intensity. Sequencing then enables determination of the relative abundance of each variant present in the sorted cells compared to the input library, which will correlate with fluorescence intensity (Peterman and Levine, 2016). However, in the case of fluorescent biosensors, the goal is not necessarily to improve the brightness, but rather the dynamic range (i.e. the relative change in emission between the ligand-bound and ligand-free (apo) states (Δ*F/F*_0_)).

A previous study has demonstrated a sort-seq assay relying on three sequential rounds of threshold sorting in the presence and absence of the ligand, that enriches for variants based on dynamic range rather than brightness (Nadler et al., 2016). While functional biosensor variants were correctly identified, this method exhibits certain biases and limitations. For example, only variants that exhibit a direct relationship between ligand concentration and fluorescence intensity (turn-on) will be enriched, disregarding the possibility of functional variants with the opposite response (turn-off). In addition, the output metric, enrichment, only correlates with dynamic range and doesn’t account for differences in brightness necessitating downstream experiments to quantify these variables for each variant. An ideal method would directly measure the brightness in each state as well as the magnitude and direction of fluorescence change for many variants in parallel to better understand the relationships between these aspects of biosensor function and sequence.

An alternate sort-seq experimental design relies on the sorting of cells into multiple bins spanning the range of fluorescence intensity in order to infer the fluorescence distributions of different variants in an unbiased fashion (Peterman and Levine, 2016). The inferred mean intensity in the ligand-bound and ligand-free states can then be used to calculate the dynamic range for each variant. This binned sort-seq approach should be generally useful for identifying turn-on as well as turn-off sensors across the entire range of observed fluorescence intensity. Here we demonstrate the application of binned sort-seq assays to efficiently screen libraries of biosensor variants. Sort-seq characterization of a previously described maltose-binding protein (MBP) domain-insertion library (Nadler et al., 2016) validates the robustness of this method to identify high-dynamic-range variants independent of baseline brightness and direction of fluorescence change upon ligand binding. Optimization of a pyruvate biosensor (Arce-Molina et al., 2019) through linker mutagenesis screened in mammalian cell culture demonstrates the generalizability of the assay to various contexts and library designs. Machine learning models trained on the resulting data enable prediction of dynamic range from linker sequence. Insights into the biochemical features underlying biosensor function can be drawn from models to aid in further optimization efforts. Together these experiments establish the use of sort-seq assays to efficiently develop high-dynamic-range biosensors.

## Results

### Sort-Seq assay of an MBP domain-insertion library

A library of circularly permuted GFP (cpGFP) insertions into MBP (Nadler et al., 2016) was used as a test-case for the sort-seq screening of biosensor libraries in *E. coli*. The transposon-based cloning strategy used to construct this library simplifies a common first step in biosensor design, the identification of a suitable insertion site in the LBD. This method relies on the random insertion of a modified transposon sequence throughout the LBD which is subsequently replaced with the cpGFP sequence through Golden Gate assembly (Fig. 1A). This library has previously been characterized by an enrichment assay that identified high-dynamic-range variants (MBP-169 and MBP-170) that serve as positive controls (Nadler et al., 2016). An initial enrichment sort was performed to select for fluorescent variants above a threshold defined by non-fluorescent cells. This sort removes the non-productive (out-of-frame and reversed insertions generated by the transposon method) as well as in-frame insertions that result in improper folding of the inserted cpGFP. Enrichment scores (*E* = *log*_2_(*f_sort_/f_naive_*)) calculated from read counts of variant sequences in the naive and sorted libraries revealed enrichment of in-frame variants (p < 0.001, Mann-Whitney U test, Data S1), and overall consistency with the results (Nadler et al., 2016) (Spearman’s ρ = 0.6, Fig. S1). These results confirm the usefulness of the transposon cloning strategy for generating many domain-insertion variants in parallel and that a single threshold sort can selectively enrich the desired fluorescent variants.

**Figure 1.**
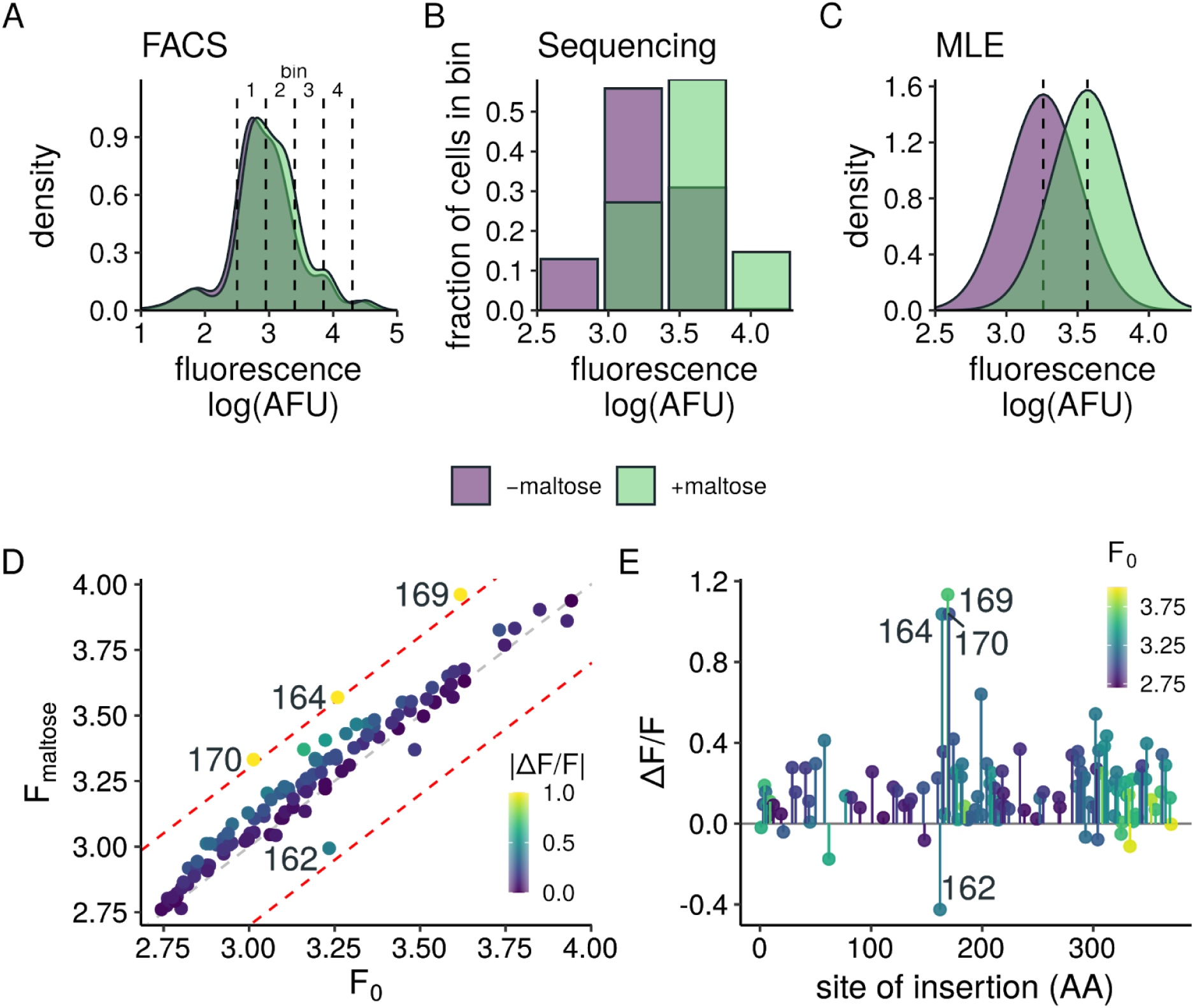
Sort-seq assay of an MBP domain-insertion library. **(A)** Fluorescence distributions for the MBP domain-insertion library ±1mM maltose. Cells were sorted into the 4 bins indicated by dashed lines. **(B)** Sequencing reads are used to estimate the relative fraction of cells containing a given variant sorted into each bin. **(C)** Maximum likelihood estimation is used to infer the log-normal distribution parameters μ and σ represented by shaded regions. Dashed lines represent the estimated log-mean (σ) for each condition. Panels B and C represent the sequencing data and MLE distributions for variant MBP-164. **(D)** Estimates of the mean fluorescence intensity for 113 MBP domain-insertion variants in the presence (F_maltose_ and absence (F_0_) of 1mM maltose. Colours represent absolute value of ΔF/F. Red and grey dashed lines indicate |ΔF/F|=1.0 and 0.0 respectively. **(E)** Profile of ΔF/F estimates for insertions along the MBP primary sequence. Colours indicate the baseline brightness for each insertion-site.

Estimates of the dynamic range for variants in the MBP library were obtained by binned sort-seq. *E. coli* expressing the enriched library were sorted into 4 equal width bins spanning the range of log fluorescence of the entire library (Fig. 1A). Cells were sorted in two batches to obtain fluorescence intensity measurements in the presence or absence of maltose (Fig. 1B). Read counts of variant sequences in each bin were used to estimate the mean and standard deviation parameters assuming a log-normal distribution (Fig. 1C). In all, estimates of brightness and dynamic range for 113 variants out of a possible 370 are present after filtering out lowly represented sequences (Fig. 1D, Data S2). These data provide a thorough examination of the potential for biosensor function of many insertion sites across the protein.

A majority (93/113 variants) exhibited little to no detectable change in fluorescence (|ΔF/F| < 0.3) illustrating the relative rarity of allosterically coupled variants. Positive controls MBP-169 (ΔF/F = 1.13) and MBP-170 (ΔF/F = 1.04) exhibited the largest maltose induced changes, and insertion at amino acid 164 exhibited a response of similar magnitude (ΔF/F = 1.04). In contrast, insertion at MBP-162 produced a biosensor with the opposite response, exhibiting a decrease in fluorescence with the addition of maltose (ΔF/F = −0.43). The proximity of the highest functioning variants in the protein sequence supports the notion that ligand binding is not necessarily allosterically coupled to specific residues but instead broader regions of the protein surface (Fig. 1E). Construction and testing of MBP-162 and MBP-164 in a 96-well plate fluorescence assay confirmed the negative and positive responses to maltose estimated by sort-seq (Fig. S2). These two variants were not detected in the previously published screen likely because of low abundance in the initial library due to intrinsic bias of the transposon reaction, each comprising approximately 1 in 10^5^ sequences. Such large disparities in variant abundance are common in many massive mutagenesis libraries. The ability to estimate dynamic range of low abundance variants highlights the potential for this assay to expand to even more diverse libraries.

### Sort-seq assay of a pyruvate biosensor linker library

We next asked if the sort-seq assay could be used to optimize the linker regions of an existing SFPB in mammalian cells. Previous studies of various SFPBs show that increases in dynamic range can be achieved by altering the linkers connecting the LBD and circularly permuted fluorescent protein (Marvin et al., 2011; Nadler et al., 2016; Nakai et al., 2001). We focused on the pyruvate biosensor, PyronicSF, since it functions robustly in mammalian cells. PyronicSF consists of cpGFP inserted into the pyruvate sensitive transcription factor PdhR (Arce-Molina et al., 2020). The linker and FP sequences of PyronicSF (Leu^1^Glu^2^-cpGFP-Thr^3^Arg^4^, referred to as L^1^ E^2^-T^3^R^4^) are derived from the well-characterized SFBP GCaMP3 (Tian et al., 2009) and were not optimized for PyronicSF (Fig. 2A). We hypothesized that tailoring the linkers to the insertion context would yield improvements to the dynamic range. A library consisting of 2,304 unique linker combinations was generated by substituting residues in linker 1 (L^1^E^2^) and 2 (T^3^R^4^) with amino acids from the sets {A, E, G, L, P, Q, R, V} and {A, G, P, R, S, T} respectively (Fig. 2A). This degenerate library encodes the original linker set, which is useful as a positive control, while also sampling from amino acids with diverse biochemical features. The linker library was transfected into the HEK293T Landing Pad line (Matreyek et al., 2017) and sorted to enrich for recombined cells ensuring genomic integration of a single variant per cell. Recombined cells were sorted into equal-width bins spanning the range of log fluorescence intensity in the presence or absence of 10mM pyruvate (Fig 2B).

**Figure 2.**
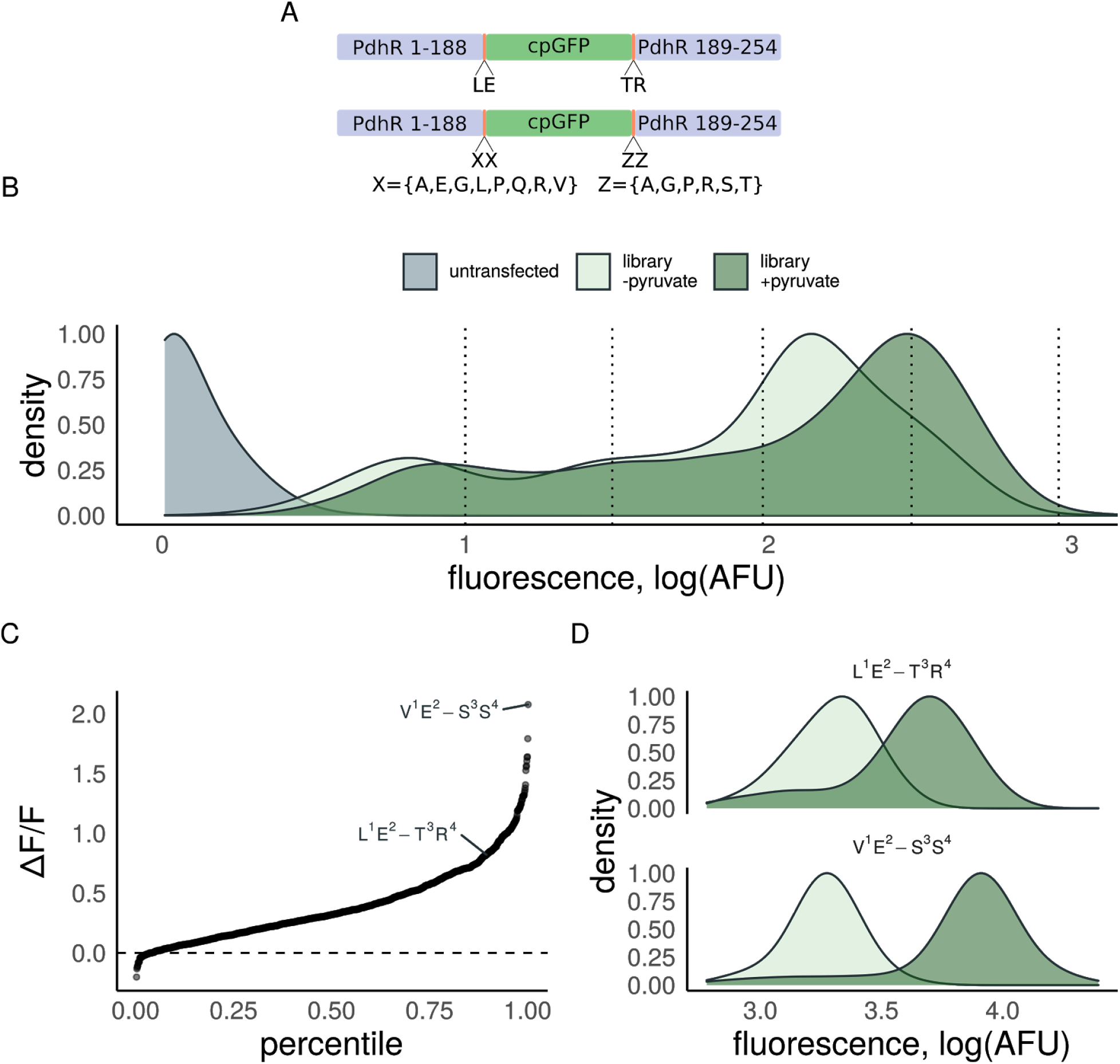
Sort-seq assay of a PyronicSF linker library. **(A)** Schematic depicting the domain organization of PyronicSF (top) and the designed linker library (bottom) **(B)** Fluorescence distributions for the PyronicSF linker library expressed in HEK293T Landing Pad cells with (dark green) or without (light green) 10mM pyruvate. Cells were sorted into the 4 bins indicated by dashed lines. **(C)** Dynamic-range estimates for 1,023 PyronicSF linker variants ordered from lowest to highest ΔF/F. The original linkers (L^1^E^2^-T^3^R^4^) and top variant (V^1^E^2^-S^3^S^4^) are labelled. (D) Flow cytometry fluorescence distributions for HEK293T cells expressing either PyronicSF or the top variant identified from sort-seq assay with (dark green) or without (light green) 20mM pyruvate. Dynamic-range is increased from ΔF/F = 1.15 for the original linkers to ΔF/F = 2.99 for variant V^1^E^2^-S^3^S^4^ validating the estimates from sort-seq assay.

Sort-seq estimates of dynamic range were obtained for 1,023 unique linker combinations (Fig. 2B, Data S3). Similar to the domain-insertion library, variation in the linker regions produced changes in brightness spanning over an order of magnitude. The estimated dynamic range was also found to be highly variable with values ranging from ΔF/F = −0.20 to 2.08 (Fig. 2C). The PyronicSF parent sequence, L^1^E^2^-T^3^R^4^, served as an internal positive control and was found to have a dynamic range (ΔF/F = 0.82) comparable to the published *in vitro* value (ΔF/F = 1.5) (Arce-Molina et al., 2020).

The variant with the highest estimated dynamic-range, V^1^E^2^-S^3^S^4^ (ΔF/F = 2.08) was constructed and tested by flow cytometry alongside the original L^1^E^2^-T^3^R^4^ linkers confirming the increase in dynamic-range observed in the sort-seq measurements (Fig. 2D). Most single substitutions to the parent sequence (21/24) resulted in a decrease in dynamic range with the substitution of Leu^1^ with Val^1^ producing the largest increase in function (Fig. S3). As would be expected given the proximity of the amino acids within a given linker as well as the presumed physical proximity of the two linkers, the effect of multiple substitutions results in epistatic interactions. In the case of V^1^E^2^-S^3^S^4^, the measured dynamic range is much greater than the linear sum of the effects of the comprising single substitutions (Fig. S3; ΔF/F = 2.08 compared to ΔF/F = 0.57 assuming additivity). Testing substitution effects at each position in serial, as is common in directed evolution experiments (Reetz and Carballeira, 2007), would not arrive at this combination evidencing the utility of testing combinatorial libraries.

In addition to discovering highly functional variants, another benefit of this approach is the opportunity to learn from the numerous suboptimal variants. Machine learning algorithms trained to predict functional activity from protein sequence can assist in elucidating the biochemical determinants of function and predict additional sequences to test (Alley et al., 2019; Bedbrook et al., 2019; Biswas et al., 2020; Wu et al., 2019; Xu et al., 2020; Yang et al., 2018). To this end, the PyronicSF linker sequences were encoded as numerical vectors using the VHSE amino acid descriptor (8 principal components score vectors derived from hydrophobic, steric, and electronic properties) (Mei et al., 2005). These biochemical encodings of linker sequences were then used as features to train a random forest regression model (Breiman, 2001; Probst et al., 2019; Svetnik et al., 2003) to predict sort-seq derived ΔF/F values. The model was trained on an 80% split of the dataset and prediction performance was tested on the held out 20%. Predictions of test set function correlated with measured dynamic range (R^2^ = 0.64, Fig. 3, Data S4) suggesting strong predictive power of the model.

**Figure 3.**
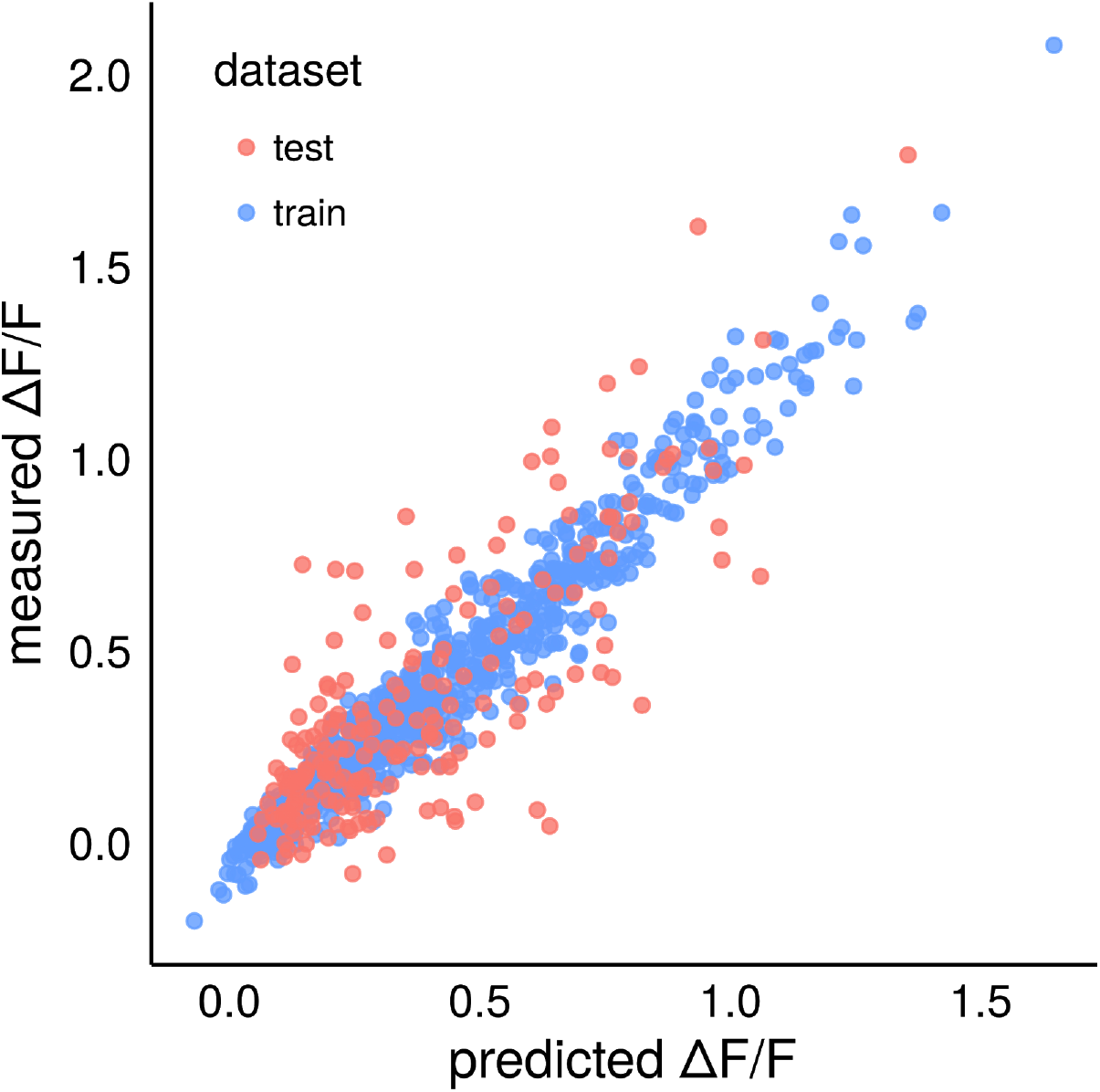
Dynamic-range can be predicted from linker sequence. PyronicSF linker variants were numerically encoded using VHSE biochemical descriptors. A random forest model was trained using an 80% split of the encoded linker data (training set, blue) to predict dynamic-range. Model performance was evaluated on the held-out 20% (test set, red). A strong correlation (R^2^ = 0.64) was observed between model predictions and measured ΔF/F.

Insights into the sequence determinants of biosensor function can be drawn from interpreting how useful the various biochemical features are for predicting function. The importance of a given feature can be estimated from the change in model accuracy when predictions are made using a permutation of that feature vector (Breiman, 2001). For example, the most important feature according to this method describes electronic properties (VHSE_5_) of the second amino acid in the N-terminal linker (E^2^ in L^1^E^2^-T^3^R^4^ and V^1^E^2^-S^3^S^4^; Fig. 4A). Variants containing glutamic acid at this position exhibited the highest mean dynamic range (ΔF/F = 0.66 ± 0.42; Fig. 4B). Interestingly, glutamic acid at this position is featured in many of the GCaMP sensor designs (GCaMP2, GCaMP3, and GCaMP6M) in which detailed structural (Akerboom et al., 2012; Chen et al., 2013) and biochemical (Barnett et al., 2017) study have found the charged side chain to preferentially stabilize the protonated state of the fluorophore in the ligand-bound conformation which underlies the fluorescence switching mechanism. In contrast, substituting positively charged arginine at this position resulted in the lowest mean dynamic range (ΔF/F = 0.20 ± 0.16; Fig. 4B). The second most important feature corresponds to backbone properties (VHSE_8_) of the first amino acid. This feature segregates proline, the most deleterious amino acid at this position (ΔF/F = 0.12 ± 0.12, Fig. 4C), from the other amino acids due to its unique cyclic structure. In contrast, variants containing the less constrained small hydrophobic amino acids alanine (ΔF/F = 0.53 ± 0.37, Fig. 4C) and valine (ΔF/F = 0.52 ± 0.39, Fig. 4C) exhibited the highest mean dynamic range. Despite the presumed importance of glutamic acid at amino acid 2, the N-terminal linker combination with the highest mean dynamic range was A^1^A^2^ (ΔF/F = 1.02± 0.39, Fig. 4D) followed by V^1^E^2^ (ΔF/F = 0.95 ± 0.49, Fig. 4D). While the features of the C-terminal linker were found to be less important for prediction accuracy (Fig. 4A), this is not to say that these positions do not contribute to biosensor function. Rather, C-terminal substitution effects can be substantial and seemingly more dependent on interactions with the other linker amino acids (Fig. S4).

**Figure 4.**
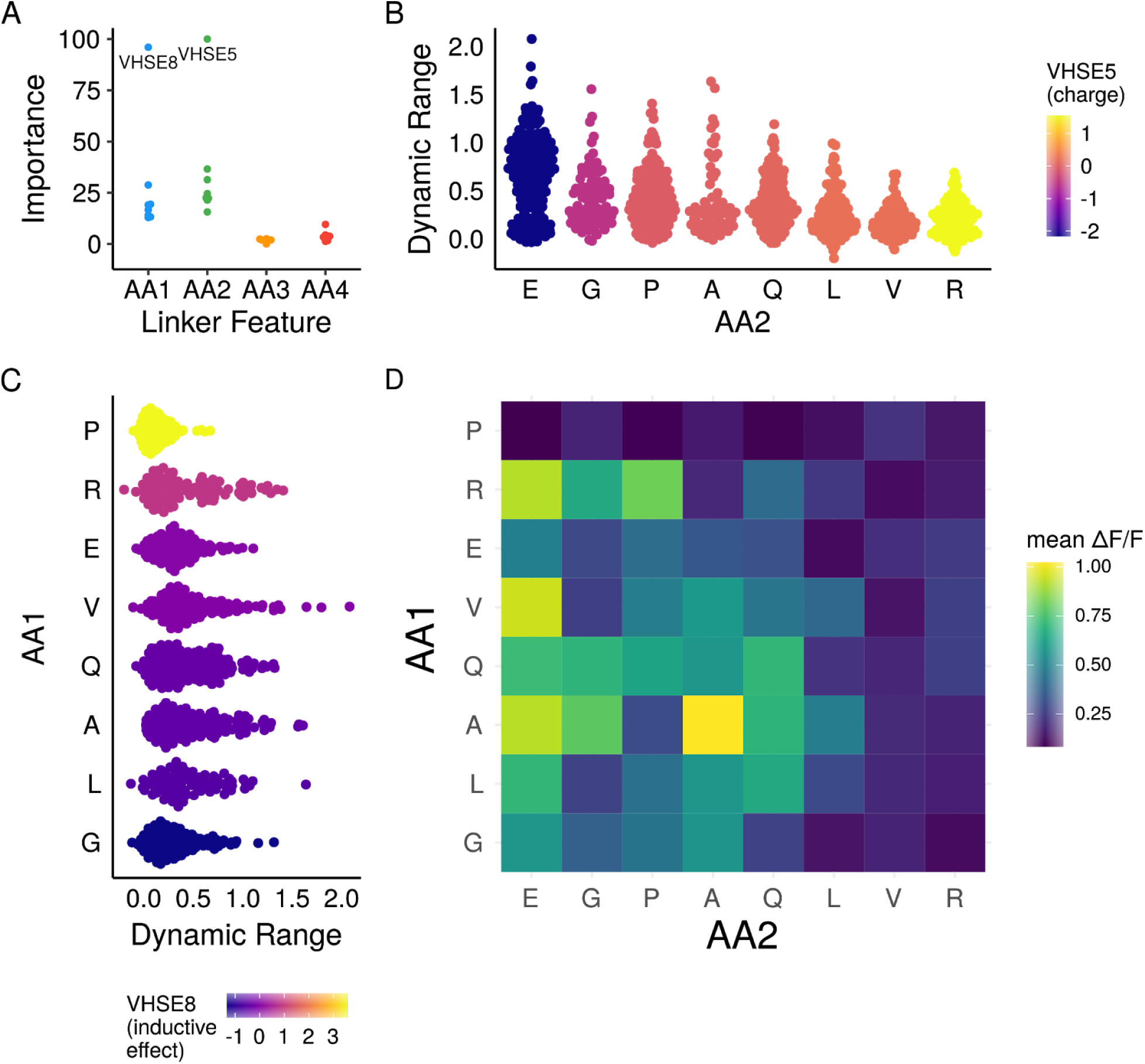
Analysis of the biochemical basis of linker function. **(A)** The importance of each feature can be estimated from the impact on model importance. Two features of the N-terminal linker amino acids, VHSE_8_ of the first and VHSE_5_ of the second amino acid are the most important. **(B)** Distributions of variant dynamic-range estimates sorted by VHSE_5_ of second N-terminal linker amino acid. Negatively charged glutamic acid exhibits the highest mean ΔF/F, while positively charged arginine exhibits the lowest. **(C)** Distributions of variant dynamic-range estimates sorted by VHSE_8_ of the first N-terminal linker amino acid. VHSE_8_ separates proline, which is distinctly deleterious, from the remaining amino acids. **(D)** Mean dynamic-range for N-terminal linker pairs highlights certain combinations that on average exhibit increased function such as A^1^ A^2^ and V^1^E^2^.

Training the model with variable training set sizes reveals sharp decreases in the mean absolute prediction error when using up to 20% of the data and diminishing returns upon further increases (Fig. S5A). This result demonstrates a machine learning model trained on a subset of a complex library is useful for efficiently prioritizing untested sequence variants to evaluate experimentally. To test this hypothesis *in silico*, the random forest model trained on 20% of the data (241 variants) was used to predict the dynamic range for all 2304 possible combinations encoded by the degenerate bases. A second library that could be encoded by degenerate bases was then designed by selecting amino acids at each position that appeared in more than 10% of the top 100 predicted variants. Comparing the distribution of dynamic range across all variants in the original library to the predicted high-dynamic-range subset reveals a shift towards increased function (Fig. S5B). By combining sort-seq assays with machine learning guided library design, we can both increase the number of variants tested as well as the likelihood a given variant will exhibit increased function.

## Discussion

Technologies to construct protein variants in parallel have outpaced the ability to assay these constructs for function. Here we show that binned sort-seq assays enable estimation of biosensor dynamic range by directly quantifying the fluorescence intensity of ligand-bound and unbound states for many variants in parallel. The application of this method to domain-insertion as well as linker variant libraries presents a high-throughput approach to the development of novel biosensor constructs as well as optimization of existing SFPBs.

Characterization of an MBP domain-insertion library using the sort-seq assay both confirms previous findings and expands the number of high-dynamic-range variants detected. Identifying functional biosensors in a pooled library is made especially difficult given that high-dynamic-range variants can exhibit differences in brightness and direction of fluorescence change. In the case of the MBP domain-insertion library, the three variants with the largest dynamic range (164, 169, 170) differ significantly in ligand-free brightness and another variant (162) exhibits a considerable dynamic range but with an inverse response to maltose. Despite these differences, these variants were all correctly identified as high-dynamic-range biosensors, highlighting the advantages of the binned sort-seq approach for discovering functional biosensors independent of brightness and response direction.

The ability to screen biosensor libraries in both live *E. coli* and HEK293T cells provides a significant advantage in regards to screening conditions. Although protein function can be sensitive to the context of expression, SFPBs are commonly developed using assays of purified protein or expressed in *E. coli*. Differences in protein translation, trafficking, and analyte abundance can lead to reduced performance when changing the expression context. Functional screens performed in mammalian cell culture using the HEK293T Landing Pad line can reduce these differences when developing sensors for *in vivo* mammalian studies (Tian et al., 2009). The assay conditions can be even further refined by converting a given cultured mammalian cell of interest into a Landing Pad line using a simple lentiviral toolkit (Matreyek et al., 2019). Developing biosensors using the combination of Landing Pad cell lines and sort-seq simplifies the engineering process by screening for biosensors in an appropriate context.

Biosensor engineering is an iterative process. Each round of mutagenesis and screening ideally will result in improvements in biosensor function as well as insights to guide future rounds of optimization. Physiologically relevant differences in concentration, such as the estimated 12μM difference between astrocyte cytosolic and mitochondrial pyruvate concentrations, are relatively small compared to the sensitive range of the biosensor (approximately 0.1mM to 10mM) (Arce-Molina et al., 2020). The 160% improvement in dynamic range resulting from our assay of PyronicSF linker variants will enable more accurate detection of such small but physiologically important changes in pyruvate concentration. In addition to the improved variants identified by this assay, the resulting dataset as a whole is a valuable resource to guide future experiments. In most biosensor engineering experiments, only the most functional variants are sequenced. However, obtaining sequence-function pairs for variants with improved function as well as mutations that decrease function is useful to better understand the sequence-function landscape. In the case of PyronicSF, mutational paths from the original linker to an improved sequence mostly traveled through intermediates with decreased function. Understanding these epistatic interactions between amino acids is essential for identifying optimized linker combinations. Machine learning algorithms applied to this problem show promise for accurately predicting dynamic range from linker sequence, identifying biochemical features contributing to biosensor function and designing new libraries biased towards high-dynamic-range variants.

In this study, variation was limited to either the site of insertion or the composition of the linkers. For a fixed insertion site such as PyronicSF, some linker variants produce strong coupling while others exhibit no coupling of ligand binding with fluorescence. This suggests that the arbitrary linkers imposed by the transposon cloning method might mask insertion sites that would be functional with different linkers. In addition, analysis of model performance revealed that only a fraction of the collected linker data for this insertion-site was required to generate robust predictions. This result suggests experimental resources could be better spent on a shallow survey of a more complex library than a comprehensive examination of all possible combinations at a few positions. Highly complex libraries containing variable linkers at each insertion site can be constructed using oligo library synthesis methods (Coyote-Maestas et al., 2020) and should be amenable to characterization by sort-seq. Studying hundreds of linker combinations at many insertion sites will help elucidate the interactions between these features.

Furthermore, the increase in sequence diversity generated by combinatorial linker and insertion-site libraries might uncover highly functional variants that would be difficult to discover by testing each parameter separately. In the short term, binned sort-seq assays provide an efficient method for systematically generating and improving SFPBs for scientific research. After many such experiments, the accumulated data might yield general insights into the principles governing biosensor design.

## Methods

### Domain-insertion library cloning

The DNA coding sequence for amino acids 1-370 of *E. coli* MBP (malE) was obtained as a gBlock (IDT DNA, Table S1). BsaI sites were added to ends by PCR with complementary overhangs for golden gate cloning into pATT-Dest (Addgene plasmid #79770) using primers MBP-BsaI-GG-F and MBP-BsaI-GG-R (the sequences for all primers used for cloning can be found in Table S1). Mu-BsaI transposon was digested from pUC-KanR-Mu-BsaI (Addgene plasmid # 79769) with BglII and HindIII in Buffer 3.1 (NEB) at 37°C overnight and purified by gel extraction (NucleoSpin Gel and PCR Clean-up). Transposition was performed using 100 ng of purified MuA-BsaI transposon, pATT-MBP plasmid DNA at a 1:2 molar ratio relative to transposon, 4 μl of 5x MuA reaction buffer, and 1 μl of 0.22 μg/μl MuA transposase (Thermo Fisher) in a total volume of 20μL. Reactions were incubated at 30°C for 18 hours, followed by 75 °C for 10 minutes. Reactions were cleaned up using the DNA Clean & Concentrator-5 Kit (Zymo Research Corp.), and eluted in 6μL of water. Transformation was performed using 2μL of reaction in 25μl of E. Cloni 10G ELITE cells (Lucigen) in 1.0-mm Bio-Rad cuvettes using a Gene Pulser Xcell Electroporation System (settings: 10 μF, 600 Ω, 1.8 kV). Cells were immediately resuspended in 975μL Recovery Media and shaken at 250rpm for 1 hour at 37°C. A 10μL aliquot of transformed cells were plated on carbenicillin (100 μg/ml) and chloramphenicol (25 μg/ml) to select for the presence of pATT plasmid backbone and transposon insertion to assess library coverage. The remaining transformed cells were pelleted and resuspended in 50mL LB with 100 μg/ml carbenicillin and 25 μg/ml chloramphenicol. Cultures were grown at 250rpm, 37°C overnight followed by plasmid DNA purification using a HiSpeed Plasmid Midi Kit (Qiagen).

Transposed pATT-MBP plasmid was digested with Esp3I and the band corresponding to MBP plus the transposon sequence was isolated by gel extraction (NucleoSpin Gel and PCR Clean-up). The isolated transposed ORF was cloned into the expression vector pTKEI-Dest (Addgene Plasmid #79784) by golden gate cloning using 40 fmoles pTKEI-Dest, 40 fmoles of purified MBP-transposon, 10 units Esp3I (NEB), 800 units T4 DNA ligase (NEB) and 1X T4 DNA Ligase Reaction Buffer (NEB) in a total volume of 20μL. The reaction was incubated 2 minutes at 37°C, 5 minutes at 16 °C for 50 cycles followed by 20 minutes at 60 °C and 20 minutes at 80 °C. Reactions were purified using a DNA Clean & Concentrator-5 Kit and eluted with 6 ml water. Transformation was performed using 2μL of reaction in 25μl of 10G ELITE E. coli (Lucigen) in 1.0-mm Biorad cuvettes using a Gene Pulser Xcell Electroporation System (settings: 10 μF, 600 Ω, 1.8 kV). Cells were immediately resuspended in 975μL Recovery Media and shaken at 250rpm for 1 hour at 37°C. A 10μL aliquot of transformed cells were plated on LB agar plates containing 50 μg/ml kanamycin and 25 μg/ml chloramphenicol to select for pTKEI-Dest backbone and transposon insertions in order to assess library coverage. The remaining transformed cells were pelleted and resuspended in 6mL LB with 1% glucose, 50 μg/ml kanamycin, and 25 μg/ml chloramphenicol. Cultures were grown at 250rpm, 37°C overnight followed by plasmid DNA purification using a QIAprep Spin Miniprep Kit (Qiagen).

BsaI sites and compatible overhangs were added by PCR amplification of cpGFP from pTKEI-Mal-B2 (Addgene Plasmid #79756) using primers cpGFP-BsaI-GG-F and cpGFP-BsaI-GG-R. Inserted transposons were replaced with cpGFP by BsaI-mediated Golden Gate cloning using 40 fmoles of pTKEI-MBP-transposon, 40 fmoles of purified cpGFP, 10 units Esp3I (NEB), 800 units T4 DNA ligase (NEB) and 1X T4 DNA Ligase Reaction Buffer (NEB) in a total volume of 20μL. The reaction was incubated 2 minutes at 37°C, 5 minutes at 16 °C for 50 cycles followed by 20 minutes at 60 °C and 20 minutes at 80 °C. Reactions were purified using a DNA Clean & Concentrator-5 Kit and eluted with 6 ml water. Transformation was performed using 1μL of reaction in 40μl of Tuner DE3 cells (Novagen) by heat shock at 42°C for 30s. Cells were immediately resuspended in 975μL Recovery Media and shaken at 250rpm for 1 hour at 37°C. A 10μL aliquot of transformed cells was plated on LB agar plates containing 50 μg/ml kanamycin and 25 μg/ml chloramphenicol to select for pTKEI-Dest backbone and transposon insertions to assess remaining transposon. The remaining culture was pelleted and resuspended in 6mL LB with 1% glucose and 25μg/mL kanamycin and grown overnight. The following day, FACS samples were inoculated from this culture and a glycerol stock was prepared for storage. Plasmid DNA was extracted from the remaining culture using a QIAprep Spin Miniprep Kit.

### MBP domain-insertion library FACS

Approximately 7 hours prior to sorting, a 100μL aliquot of the library culture to be sorted as well as FACS controls (pTKEI-MBP and pTKEI-cpGFP) were added to 5mL of MOPS EZ Rich Defined Medium (Teknova) containing 60 μg/mL kanamycin and 0.4% glycerol. Cultures were shaken at 250rpm, 37°C for approximately 2 hours until the OD600 was between 0.6 and 0.8. Expression was induced by adding 0.5mM IPTG followed by incubation for 4 hours. Samples were transported on ice to the core facility for sorting and incubated an additional 30 minutes at 37 °C, 250 rpm. Immediately before starting sort, the samples were diluted 1:100 in phosphate-buffered saline (PBS) and kept on ice.

Sorts were performed using a BD FACSAria III instrument equipped with a 488nm laser for excitation and a 530/30nm emission filter for GFP measurements and 561nm laser with 610/20 filter for mCherry measurements. To minimize the sorting of aggregated cells, E. coli expressing either GFP or mCherry were mixed evenly and the sort rate was adjusted until less than 1% of cells were double positive for green and red fluorescence. Cells expressing MBP alone were used to adjust instrument voltages and establish a baseline for cellular autofluorescence. Cells expressing the MBP-cpGFP domain-insertion library were sorted to collect cells above a threshold set at the upper range of autofluorescence. Cells were sorted into a 15mL conical tube containing 5mL LB supplemented with 1% glucose. For the initial enrichment sort, a total of 4.4*10^6^ cells were collected. Immediately after sorting, cells were incubated for 1 hour at 37°C, 250rpm. The recovered cultures were diluted to 15mL LB with addition of 1% glucose. An aliquot was diluted 1:50 and spread on LB agar plates containing 50 μg/mL kanamycin to assess cell survival. The remaining culture was incubated with 50μg/mL kanamycin overnight at 30°C, 250rpm. The next day, cultures for FACS were inoculated following the steps above and a glycerol stock was prepared to store the fluorescence enriched library. Plasmid DNA for sequencing was extracted from the remaining culture using a QIAprep Spin Miniprep Kit.

Two samples were prepared for the second round of FACS in order to perform the binned sort with and without the addition of maltose. The two samples were prepared as described above with a 1.5 hour offset to account for time taken to sort the first sample. After transporting the first induced sample to the core, 1mM maltose was added before incubating an additional 30 minutes. Immediately before starting sort, the samples were diluted 1:100 in phosphate-buffered saline (PBS) or 1:100 in PBS with added 1mM maltose and kept on ice. Cells expressing MBP or cpGFP were used to determine the range of fluorescence to be sorted. The lower bound was set at the upper range of autofluorescence indicated by MBP expressing cells. The upper bound was set to include approximately 50% of the cells expressing cpGFP. Four equal width gates on the log scale were set to span this range. The +/- maltose samples were sorted using the same gates for 1.5 hours each. The cells for each bin were collected in 5mL tubes containing 1mL LB with 1% glucose. A range of 4,200 to 444,000 cells were collected for each bin approximately proportional to the relative density of cells in each bin. Cells were recovered as described above and grown overnight. The following day, plasmid DNA for each bin was extracted using a QIAprep Spin Miniprep Kit.

### MBP domain-insertion library sequencing

The ORF to be sequenced was PCR amplified from pTKEI plasmid using primers pTKEI-seqamp-F and pTKEI-seqamp-R (the sequences for all primers used for DNA sequencing can be found in Table S1). 50μL reactions were prepared with a final concentration of 0.2ng/μL plasmid DNA, 0.25μM forward and reverse primer, 1X SeqAmp PCR buffer, 1X SeqAmp Polymerase (Clontech), 1X SYBR Green (Invitrogen). Amplification was monitored by qPCR with cycling conditions: [94°C 60s, (98°C 10s, 55°C 15s, 68°C 60s, plate read) x 29 cycles]. The number of cycles was determined such that reactions were in the exponential phase of amplification upon completion of the program. Reactions were cleaned with Ampure XP beads and eluted in 40μL elution buffer (5 mM Tris/HCl, pH 8.5).

Amplicons were fragmented and tagged (tagmented) in a 25μL reaction containing 1ng/μL amplicon, 1X TD buffer and 0.5μL TDE1 enzyme from the Nextera DNA Sample Prep Kit (Illumina). Tagmentation reactions were cleaned up using a Nucleospin column and eluted in 15μL elution buffer. Tagmented DNA was amplified with primer i5-Nex2p and a unique indexed primer per sample (i7-TXX-NEX2p). 25μL reactions were prepared containing 1μL tagmented DNA, 0.5μM forward and reverse primer, 1X KAPA HiFi Hotstart Readymix (KHF), and 1X SYBR Green. Amplification was monitored by qPCR with cycling conditions: [72°C 3 minutes, 95°C 20 seconds, (98°C 20 seconds, 52°C 15 seconds, 72°C 30 seconds, plate read, 72°C 8 seconds) x 20 cycles]. Reactions were removed during the exponential phase of amplification. PCR products were run on a 1.5% agarose gel to visualize distribution of tagmented DNA size and to estimate relative concentrations using FIJI gel analysis. Indexed samples were pooled normalizing for relative concentration. Pooled products were run on a 1.5% agarose gel, cutting out a band at approximately 500bp which was then purified using the NucleoSpin Gel and PCR Clean-up column. The concentration of the pooled library was quantified using a Qubit fluorometer and size distribution was assessed using a HS DNA chip on the Bioanalyzer 2100 instrument (Agilent). The library was sequenced using 2×75bp paired-end reads on an Illumina MiSeq (v3 Reagent kit).

### MBP domain-insertion library enrichment analysis

Paired end reads were merged using BBMerge (Bushnell et al., 2017). MBP-cpGFP insertion sites were counted using the dipseq analysis pipeline developed by the Savage lab available at (https://github.com/SavageLab/dipseq) (Nadler et al., 2016). This python package identifies reads that contain sequences originating from both MBP and cpGFP. Junction reads are then trimmed of the cpGFP sequence plus transposon scar before aligning the remaining sequence to MBP to identify the site of insertion. The output is a file containing counts for each insertion site (including out-of-frame and reverse insertions) in each sample which was used for enrichment and sort-seq analysis.

Read counts deriving from the sorted and naive libraries were converted to fractional counts: 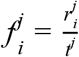 where *r* is the number of read counts for insertion at position *i* in sample *j* and *t* is the total number of reads in sample *j*. Variants with a count of 0 in any sample were filtered out and not used in further analysis. Enrichment scores were calculated as 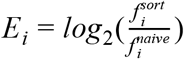.

### Maltose biosensor plate reader assay

MBP-162 and MBP-164 were cloned by Gibson assembly and tested for function individually (sequences for biosensor constructs can be found in the Supplementary Text). The backbone was opened up by PCR amplification of pTKEI-MBP using primers MBP-162-GA-F1/R1 and MBP-164-GA-F1/R1. The cpGFP insert was amplified from pTKEI-cpGFP using primer sets MBP-162-GA-F2/R2 and MBP-164-GA-F2/R2. 20μL Gibson assembly reactions were prepared with 50ng backbone, 5:1 molar ratio of insert to backbone and 1X Gibson Assembly Master Mix (NEB) and incubated at 50°C for 1 hour before transformation into One Shot TOP10 Chemically Competent *E. coli* (Invitrogen). Single colonies were picked from LB agar plates with 50μg/mL kanamycin the next day and Sanger sequenced to confirm correct assembly. Expression plasmids with the correct sequence were transformed into Tuner DE3 cells for testing. Single colonies of each biosensor variant to be tested were picked into a deep 96-well plate with each well containing 200 mL of LB supplemented with 10% glycerol, 1% glucose and 60 μg/ml kanamycin. Six wells were inoculated for each variant to accomodate 3 replicates for 2 conditions (+/- ligand). Plates were incubated overnight at 30°C, 300rpm. The following day, 5μL of culture was used to inoculate 200μl MOPS EZ Rich Defined Medium (Teknova) containing 60 μg/mL kanamycin and 0.4% glycerol in a clear bottom black 96-well plate. The plate was incubated at 37°C, 300rpm in a Molecular Devices i3 plate reader while measuring OD600 and fluorescence (485/5nm excitation and 515/5 nm emission) every 600s for 2.5 hours. Upon OD600 exceeding 0.6 for all wells, expression was induced by adding 0.5mM IPTG. Following 2 hours of incubation to allow for adequate protein expression, either 1mM maltose in PBS or equal volume of PBS alone was added to each well. After adding ligand, wells were incubated and monitored for another 1 hour. Dynamic range was calculated as Δ*F/F*_0_ = (*F_t_* − *F*_0_) / *F*_0_ where *F_t_* is the fluorescence at time *t* and *F_0_* is the fluorescence at the start of experiment. The difference between the calculated Δ*F/F_0_* for each variant with and without maltose was calculated to account for changes in fluorescence due to increased protein expression over time.

### PyronicSF linker library cloning

DNA encoding PyronicSF was obtained from Addgene (Plasmid #124812) in the expression vector pcDNA3.1(-). PyronicSF was cloned into the holding plasmid HC_Kan_RFP-p7 (Addgene Plasmid #100615) by Golden Gate Assembly. PCR was used to amplify the coding region in 4 parts to add BsaI recognition sites and compatible overhangs to ends while also removing internal BsaI/Esp3I sites that would interfere with future cloning steps using primers PyronicSF-GG-comp-F1/2/3/4 and PyronicSF-GG-comp-R1/2/3/4. Assembly was performed in a 10μL reaction containing 13 pmol of each PCR fragment and holding plasmid, 10 units BsaI-HFv2 (NEB), 100 units T4 DNA ligase (NEB), and 1X T4 DNA ligase buffer (NEB). The reaction was incubated at 37°C for 5 minutes, 16°C for 10 minutes for 40 cycles, followed by 16°C for 20 minutes, 60°C for 30 minutes and 75°C for 6 minutes before transformation into One Shot TOP10 Chemically Competent *E. coli*.

Mutations to the PyronicSF linkers were generated using degenerate primer PCR. The 5’ linker was mutated using two SNA codons and the 3’ linker using two VST codons. The backbone sequence including PdhR was opened by amplification with primers PyronicSF-VSTx2-GA-F1 and PyronicSF-SNAx2-GA-R1. cpGFP was amplified using primers PyronicSF-VSTx2-GA-F2 and PyronicSF-SNAx2-GA-R2. Overlapping homologous sequences were annealed by Gibson assembly. The reaction was cleaned using a DNA Clean & Concentrator-5 Kit (Zymo Research Corp.), and eluted in 6μL of water. The library was transformed by electroporation using 1μL of cleaned reaction in 25μL E. cloni 10G ELITE cells. A 5μL aliquot of recovered transformation was plated on LB agar containing 50μg/mL kanamycin. The remaining recovered culture was diluted into 50mL LB containing 50μg/mL kanamycin and grown overnight at 37°C, 250rpm. Transformation produced well over 20,000 unique transformants, enough to cover 2,304 variants in library 10X.

A golden gate compatible recombination plasmid, referred to as EMMA-attB-Dest, containing a Bxb1 attB site was cloned using parts from the Extensible Mammalian Modular Assembly (EMMA) Toolkit (Martella et al., 2017). The PyronicSF linker library was cloned into EMMA-attB-Dest using an Esp3I mediated Golden Gate reaction. In a 10μL reaction containing 13pmol of the PyronicSF linker library in the holding plasmid, 13pmol of EMMA-attB-Dest, 5 units Esp3I (NEB), 100 units T4 DNA ligase (NEB), and 1X T4 DNA ligase buffer (NEB). The reaction was incubated at 37°C for 5 minutes, 16°C for 10 minutes for 40 cycles, followed by 16°C for 20 minutes, 60°C for 30 minutes and 75°C for 6 minutes. The Golden Gate reaction was cleaned using a DNA Clean & Concentrator-5 Kit, eluted in 6μL of water and transformed by electroporation using 1μL reaction added to 25μL of E. cloni 10G ELITE cells. Recovered cells were diluted into 50mL LB with 100μg/ml carbenicillin and grown overnight at 37°C, 250rpm. Plasmid DNA for transfection was purified using a Qiagen Plasmid Maxi Kit.

### HEK293T Landing Pad transfection

HEK293T Landing Pad (TetBxb1BFP) cells obtained from the Fowler lab (Matreyek et al., 2017) were cultured in Dulbecco’s modified Eagle’s medium (DMEM) containing 25mM glucose and 4mM L-glutamine (Gibco 11965092) supplemented with 10% fetal bovine serum (FBS) and 25mM HEPES. Cultures were maintained with 2μg/ml doxycycline (Sigma-Aldrich) which was removed 1 day prior to transfection. For each transfection, a total 3μg of plasmid DNA and 6μL FuGENE 6 (Promega) were combined in 300μL Opti-MEM (Gibco). A 50:50 mixture by mass of pCAG-NLS-HA-Bxb1 (Addgene Plasmid #51271), for Bxb1 recombinase expression, and a recombination plasmid containing the attB sequence were used. The DNA transfection reagent mixture was added to a 6-well plate containing 1×10^6^ freshly seeded cells per well. Cells were incubated with transfection reagents for two days before expanding each well to a 15cm plate. After 1 day of growth in the 15cm plate, 2μg/ml doxycycline was added to induce expression. Cells were grown for an additional 7 days after induction before any cytometric analysis or FACS was performed.

### PyronicSF linker library FACS

Cells at 50% confluency in a 15cm plate were detached by trypsinization, pelleted and resuspended in 2mL DMEM supplemented with 50μg/mL gentamicin. Sorts were performed using a BD Influx instrument equipped with a 488nm laser for excitation and a 530/40nm emission filter for GFP measurements and 405nm laser with 460/50 filter for mTagBFP measurements. Cells transfected with Bxb1 recombinase but no attB plasmid were used to adjust instrument voltages and establish a baseline of autofluorescence in the green channel and presence of mTagBFP expression. Cells transfected with an attB plasmid but without expression of Bxb1 recombinase were used to determine the level of green fluorescence derived from plasmid expression as opposed to genomic integration, doxycycline induced expression. Cells transfected with the PyronicSF linker library and Bxb1 recombinase were sorted to collect cells positive for GFP and negative for mTagBFP fluorescence to enrich for recombined cells. A total of 300,000 GFP+/BFP-cells were collected in a 15cm tube containing 5mL DMEM supplemented with 10% FBS and 50μg/mL gentamicin. Sorted cells were pelleted, resuspended in 800μL media and plated in a 12-well plate. Cells were expanded over about 7 days to reach 50% confluence in two 15cm plates before the second round of sorting.

The two samples for the second round of FACS were detached from 15cm plates by trypsinization, pelleted and resuspended in 2mL DMEM supplemented with 50μg/mL gentamicin and either 10mM pyruvate or an equal volume of water. Cells transfected with Bxb1 recombinase but no attB plasmid were used to adjust instrument voltages and establish a baseline of autofluorescence in the green channel. Four equal width gates on the log scale were set to span the range of log(AFU) from 1.0 to 3.0. The +/- pyruvate samples were sorted using the same gates for a duration of 1.5 hours each. The cells for each bin were collected in 5mL tubes containing 1mL DMEM supplemented with 10% FBS and 50μg/mL gentamicin. A range of 200,000 to 1,315,000 cells were collected for each bin approximately proportional to the relative density of cells in each bin. Sorted cells for each bin were individually pelleted, resuspended and plated in either a 24-well (<500,000 cells), 12-well (>500,000 and <900,000 cells) or 6-well plate (>900,000 cells). All samples were expanded to 50% in a 10cm plate before harvesting and storing at −20°C after pelleting and washing with PBS.

### PyronicSF linker library sequencing

Genomic DNA was extracted from approximately 5M cells using a Qiagen DNeasy Blood & Tissue Kit. The linker regions flanking cpGFP were PCR amplified from the genome using primers PyronicSF-LLseq-F and PyronicSF-LLseq-R.

Four replicate 50μL reactions were prepared for each sample with a final concentration of 5ng/μL genomic DNA, 0.25μM forward and reverse primer, 1X SeqAmp PCR buffer, 1X SeqAmp Polymerase (Clontech), 1X SYBR Green (Invitrogen). Amplification was monitored by qPCR with cycling conditions: [94°C 60s, (98°C 10s, 55°C 15s, 68°C 60s, plate read) x 26 cycles]. The number of cycles was determined such that reactions were in the exponential phase of amplification upon completion of the program. Replicate reactions were pooled and cleaned with a Nucleospin column and eluted in 15μL elution buffer (5 mM Tris/HCl, pH 8.5).

Pooled first round PCR products were amplified a second time with primer i5-IPE2p and a unique indexed primer per sample (i7-iPE2p-XX). 25μL reactions were prepared containing 1μL round 1 DNA, 0.5μM forward and reverse primer, 1X KAPA HiFi Hotstart Readymix (KHF), and 1X SYBR Green. Amplification was monitored by qPCR with cycling conditions: [95°C 3 minutes, (98°C 20 seconds, 60°C 15 seconds, 72°C 30 seconds, plate read, 72°C 8 seconds) x 8 cycles]. Reactions were removed during the exponential phase of amplification. PCR products were run on a 1.5% agarose gel to ensure only a single band had been produced and to estimate relative concentrations using FIJI gel analysis. Indexed samples were pooled normalizing for relative concentration. Pooled products were run on a 1.5% agarose gel, cutting out a band at 933bp which was then purified using the NucleoSpin Gel and PCR Clean-up column. The concentration of the pooled library was quantified using a Qubit fluorometer and size distribution was assessed using a HS DNA chip on the Bioanalyzer 2100 instrument (Agilent). The library was sequenced using 2×75bp paired-end reads on an Illumina MiSeq (v3 Reagent kit) loaded at a final concentration of 14pM with 15% PhiX spiked-in. Sequencing reads were processed using an R script available at https://github.com/jnkoberstein/biosensor-sort-seq. Briefly, reads were trimmed to remove all but the linker sequence using Cutadapt (Martin, 2011) and allowed degenerate codons were counted using the ShortRead R/Bioconductor package (Morgan et al., 2009). The resulting table of read counts for each linker sequence in each sample was used as input for sort-seq analysis. A relatively high amount (28.6% of reads in the naive library) of the parent sequence (including bases not allowed using the degenerate scheme) was detected likely as a result of carryover from the plasmid used as a source for cloning the linker library. These reads were included for downstream MLE fitting, but excluded from all further machine learning analyses in favor of the parent amino acid sequence encoded by allowed bases.

### Sort-seq data analysis

Sort-seq data analysis was performed as thoroughly described by Peterman *et al*. (Peterman et al., 2014; Peterman and Levine, 2016) using functions written in R available at https://github.com/jnkoberstein/biosensor-sort-seq. The sorting experiment involves *m* gates of width *w* spanning a range of logarithmic fluorescence. Sorting gate *j* is defined by its upper and lower boundaries, *L_j_* and *U_j_* Given read counts *r_ij_* the number of read counts of variant *i* in bin *j*, the mean fluorescence assuming a log-normal distribution can be estimated using a maximum likelihood approach. Raw sequencing data was processed to obtain read counts of each variant in each bin using the dipseq python package for the MBP library or a custom R script for the PyronicSF library. A proportionality constant *d_j_* relating read counts to the total number of sorted cells in each bin *b_j_* is set as

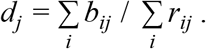

Given the count data and gate parameters, the MLE is computed as the values 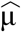 and 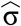 that maximize the log-likelihood function

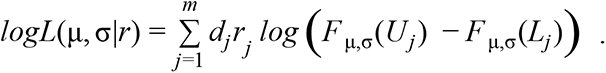

The values 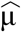 and 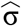 for each variant *i* were obtained by minimizing - *logL*(μ, σ|*r_ij_*) over μ and σ using the Nelder-Mead algorithm while keeping the read counts and experimental parameters fixed. The log-normal mean fluorescence is then calculated as 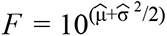. Mean fluorescence was calculated for both samples and then used to calculate dynamic range as Δ*F*/*F*_0_ = (*F_l_* − *F*_0_) / *F*_0_ where *F*_1_ is the fluorescence intensity in the sample with added ligand and *F_0_* is the fluorescence intensity in the absence of ligand. The MBP library dataset was filtered to keep only variants in which more than 100 cells were collected between the two samples and the variance in each sample was between 0.1 and 0.3. The PyronicSF library was filtered to keep only variants in which an estimate of more than 500 cells were collected for each sample and variance in each sample was between 0.1 and 0.4.

### PyronicSF-VE-SS flow cytometry

PyronicSF-VE-SS was cloned by Gibson assembly. The backbone was opened up by PCR amplification of HC_Kan_PyronicSF-p7 using primers PyronicSF-VE-SS-GA-F1/R1. The cpGFP insert was amplified from HC_Kan_PyronicSF-p7 using primer sets PyronicSF-VE-SS-GA-F2/R2. 20μL Gibson assembly reactions were prepared with 50ng backbone, 5:1 molar ratio of insert to backbone and 1X Gibson Assembly Master Mix (NEB) and incubated at 50°C for 1 hour before transformation into One Shot TOP10 Chemically Competent *E. coli* (Invitrogen). Single colonies were picked from LB agar plates with 50μg/mL kanamycin the next day and Sanger sequenced to confirm correct assembly. PyronicSF-VE-SS was cloned into EMMA-attB-Dest using an Esp3I mediated Golden Gate reaction as described for library cloning. The resulting plasmid, EMMA-attB-PyronicSF-VE-SS was transfected and recombined into Landing Pad cells. Similarly, a construct containing the original linkers, EMMA-attB-PyronicSF-LE-TR was recombined into Landing Pad cells. Recombined cells grown to approximately 50% confluency in a 6-well plate were detached by trypsinization, pelleted and resuspended in 400μL DMEM. Cytometric evaluation was performed on a BD LSR II equipped with a 488nm laser for excitation and a 530/30nm emission filter for GFP measurements and 405nm laser with 440/40 filter for mTagBFP measurements. Events were gated for live cells using FSC-A and SSC-A and single cells using FSC-A and FSC-H. Recombined cells were analyzed by gating for loss of BFP fluorescence. GFP intensity for recombined cells was analyzed in the presence and absence of 20mM pyruvate.

### Random Forest Models

PyronicSF linker sequences were converted to 32 length numerical vectors by encoding each of the 4 amino acids as the 8 representative VHSE biochemical features. Model training was performed using the R packages caret(Kuhn, 2008) with the random forest model implemented in ranger (Wright and Ziegler, 2015). An 80% split (819 variants) was used for model training and the remaining 20% (204 variants) was used for testing the final model. Fivefold cross-validation was used for hyperparameter tuning. Grid search was performed to find the optimal values for the following hyperparameters: mtry=5, 7, 13; splitrule=variance, extratrees, maxstat; min.node.size=1, 5, 7. Feature importance was calculated using the permutation method. The learning curve was generated by training the model with subsets consisting of 2.5% to 80% of the data while evaluating mean absolute error on the 20% test set and on the held out cross-validation resampling set with hyper-parameters set to mtry=7, splitrule=extratrees, and min.node.size=5.

## Acknowledgments

We thank the OHSU Flow Cytometry Core for assistance with fluorescence activated cell sorting and cytometric analysis; Yibing Jia and the OHSU Molecular Technologies Core for sequencing services; Doug Fowler and Kenneth Matrayek for sharing the HEK293T Landing Pad cell line.

---

**Figure S1.**
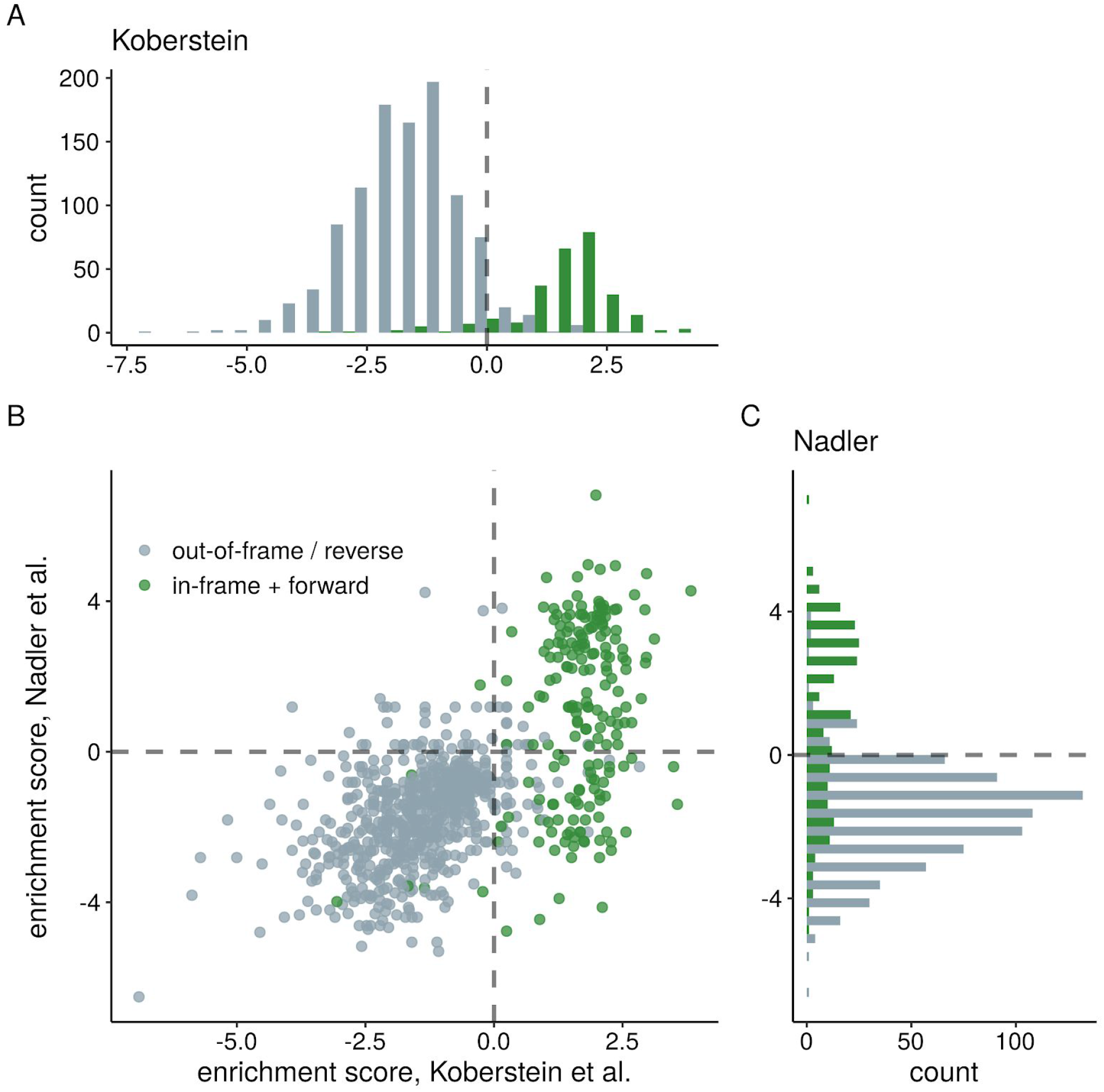
Enrichment of productive cpGFP insertions into MBP. **(A)** Distribution of enrichment values from this study for MBP domain-insertion variants following a single enrichment sort for GFP+ cells. **(B)** Comparison of enrichment values for MBP domain-insertion variants across studies reveals consistent enrichment of variants on the basis of fluorescence intensity generated by in-frame and forward insertion of GFP (Spearman’s ρ = 0.6). **(C)** Distribution of enrichment values from Nadler et al. for MBP domain-insertion variants following a single enrichment sort for GFP+ cells

**Figure S2.**
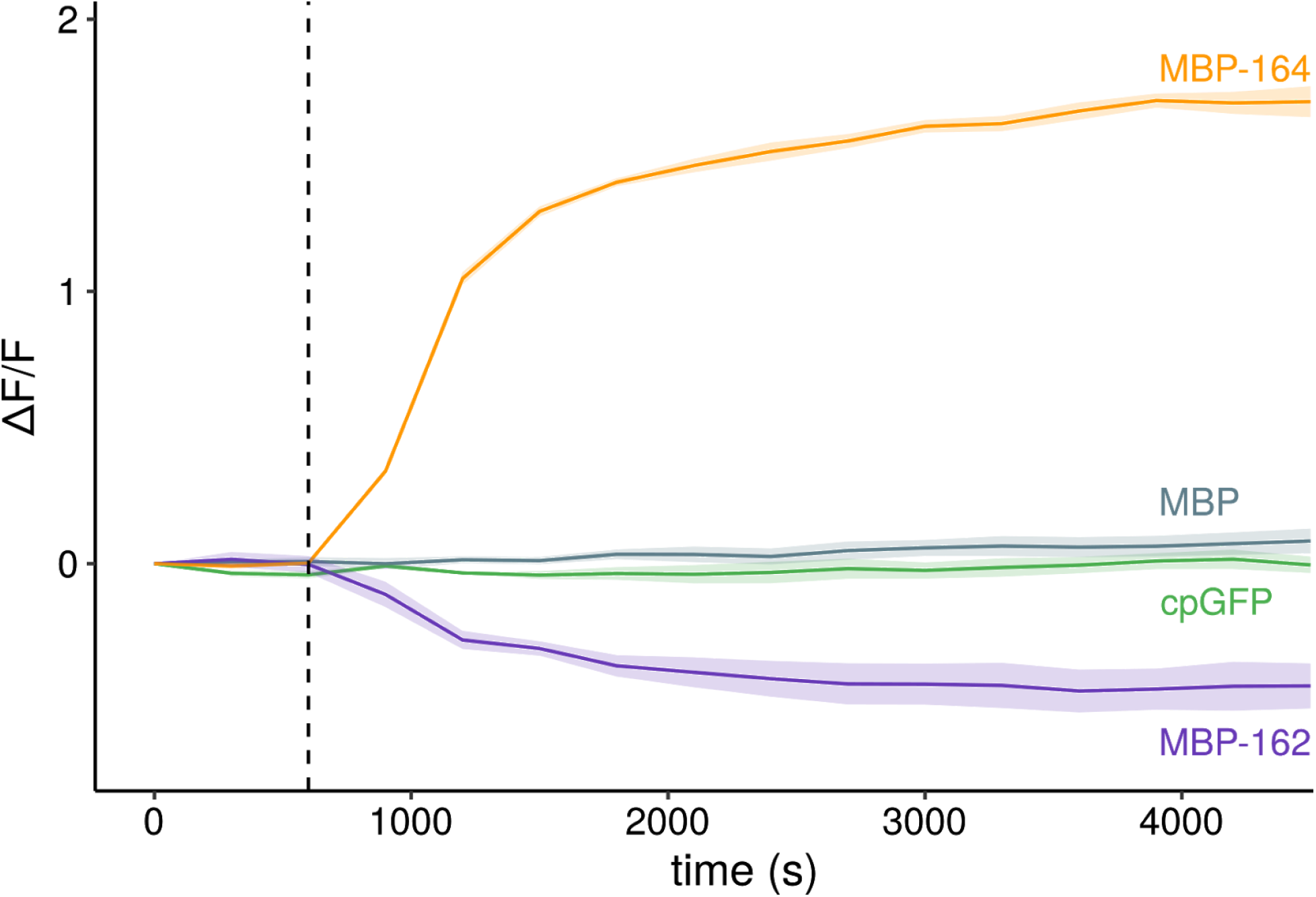
MBP-162 and MBP-164 function as biosensors with opposite responses to increased maltose concentration. The fluorescence of MBP-162 and MBP-164 were monitored following the addition of 1mM maltose (dashed line). Controls expressing either MBP or cpGFP were monitored for non-specific changes with addition of maltose. Fluorescence was measured for wells with either maltose or an equal volume of PBS added and the difference between conditions calculated to account for increases in intensity due to changes in protein expression over time. Data are mean±s.d. for three replicates for each condition.

**Figure S3.**
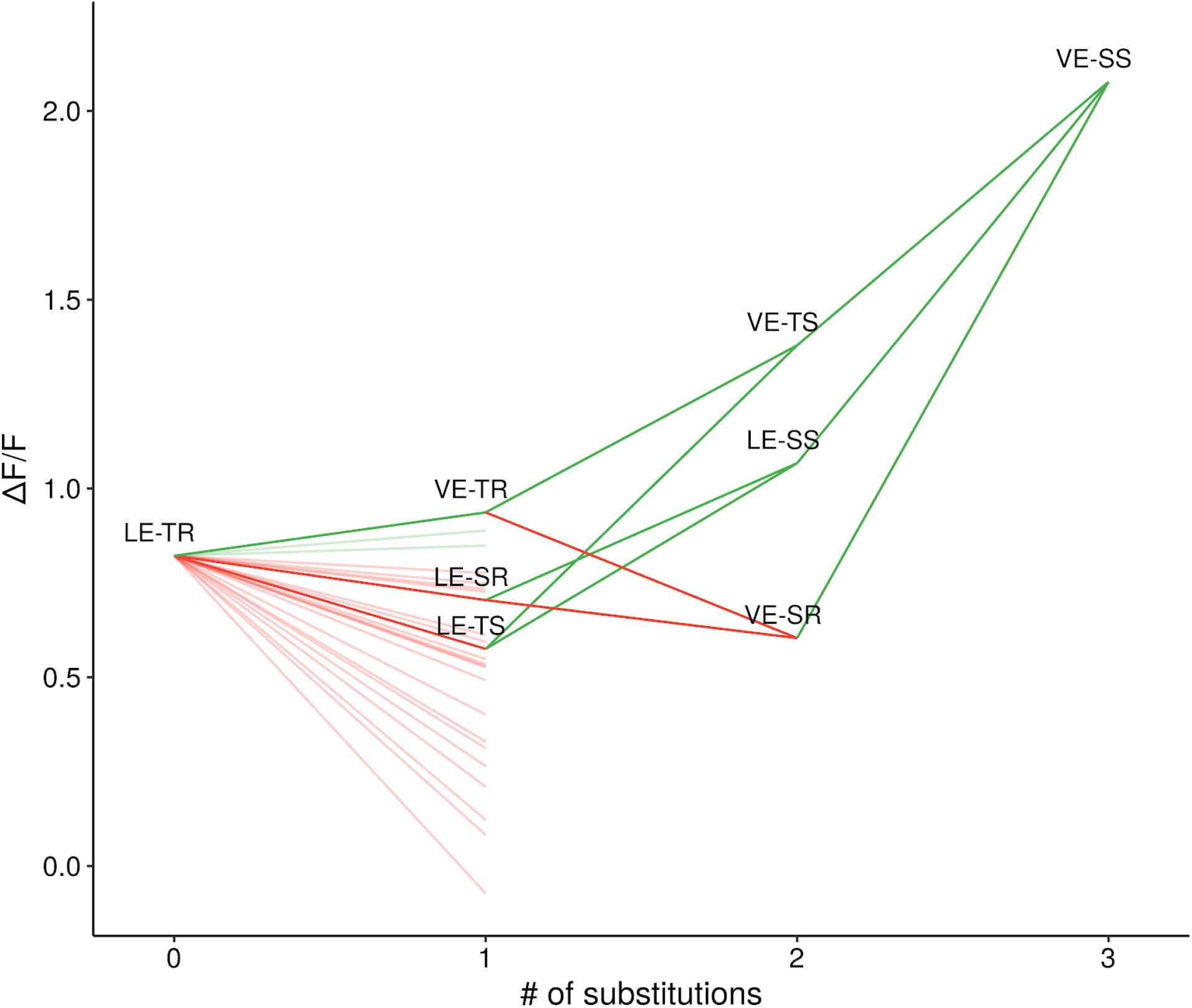
Dynamic range for linker variants versus the number of substitutions to the parent sequence. Five out of the six paths from the parent sequence (LE-TR) to the optimized sequence (VE-SS) travel through intermediates with decreased function (dark lines; green/red indicates increase/decrease in function relative to parent sequence). A majority (21/24) of the tested single substitutions to the parent sequence (additional shaded lines) result in a decrease in function. The total increase in dynamic range of VE-SS compared to the parent sequence is much greater (ΔF/F=2.08) than the sum of the effects of the comprising single substitutions (ΔF/F = 0.57).

**Figure S4.**
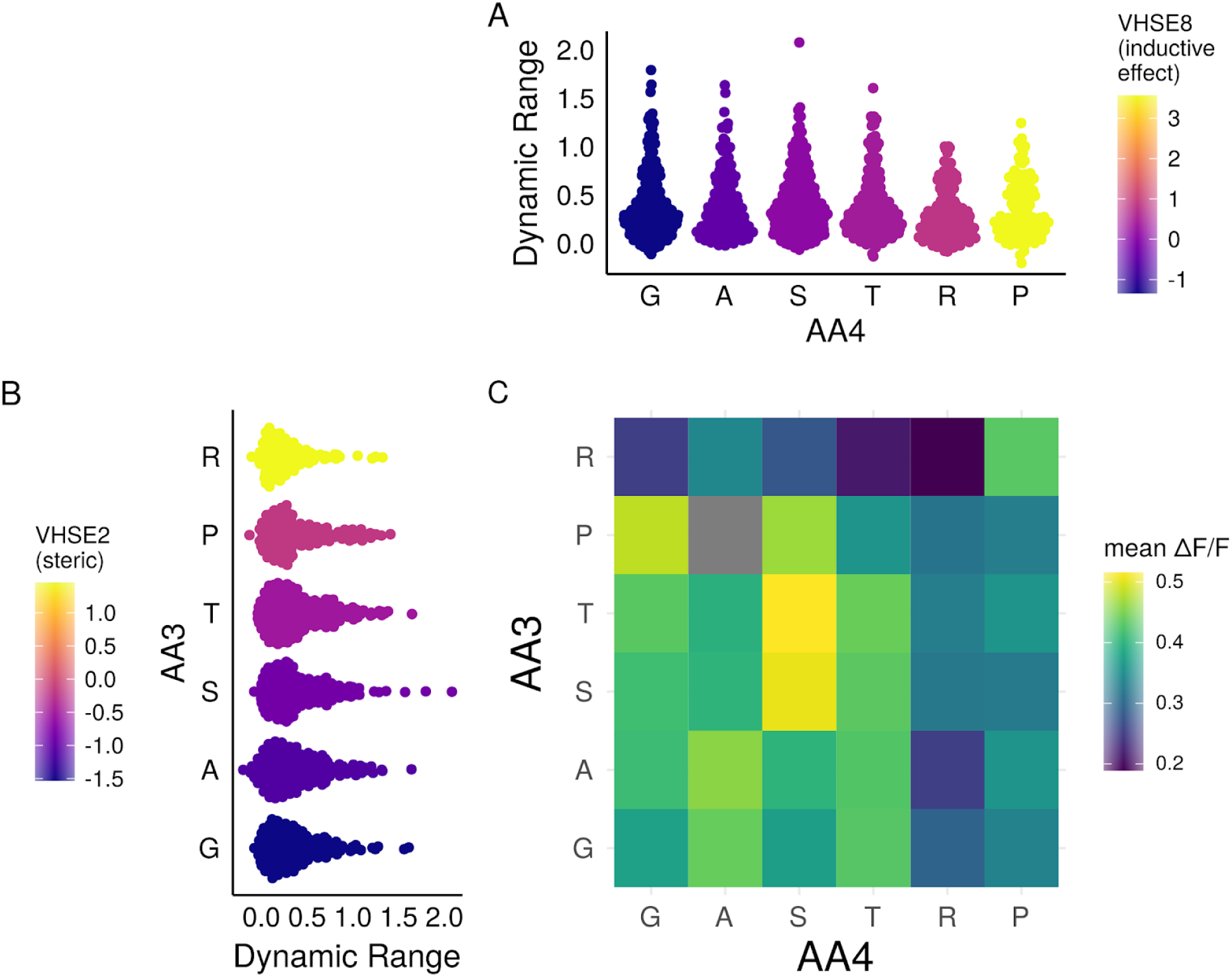
Analysis of the biochemical basis of C-terminal linker function. **(A)** Distributions of variant dynamic-range estimates sorted by VHSE_8_ of second C-terminal linker amino acid. **(B)** Distributions of variant dynamic-range estimates sorted by VHSE_2_ of the first C-terminal linker amino acid. **(C)** Mean dynamic-range for C-terminal linker pairs. The combination P^3^A^4^ is excluded (grey) as no variants with the linker combination passed filtering.

**Figure S5.**
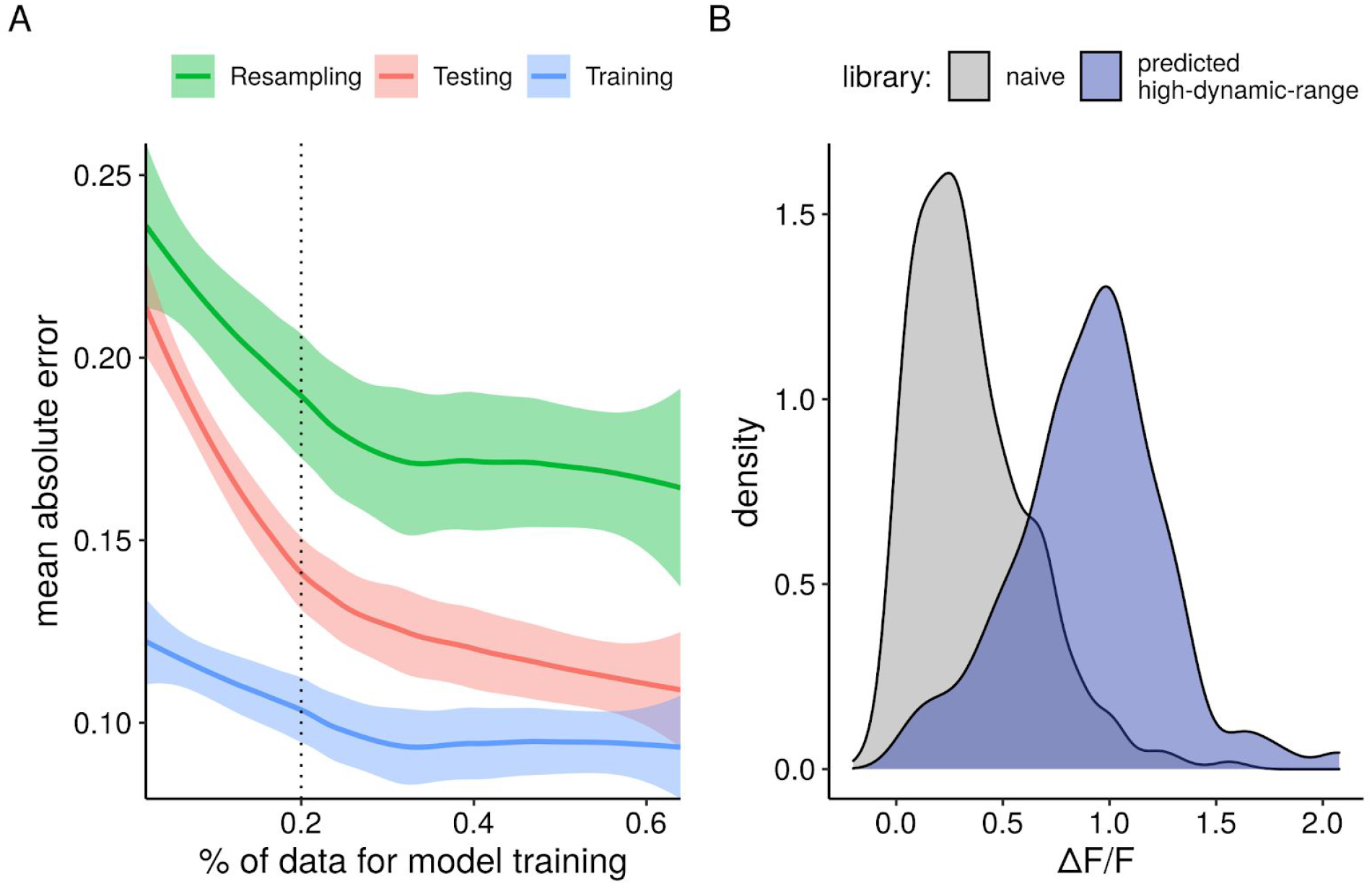
A random forest model trained on a subset of the collected data can be used to design a library with an increase in dynamic range. **(A)** Mean absolute error as a function of the percent of data used to train the model evaluated on predictions of dynamic range for the test, training, and resampled data. Lines represent a locally weighted smoothing (LOESS) fit and shaded regions represent a 95% confidence interval. The vertical dashed line represents the 20% of data used to train the model in panel B. **(B)** Distribution of dynamic range estimates for variants in the naive library compared to the high-dynamic-range variants predicted by a model trained on 20% of the data (192 variants).

