## Supplementary Text for "A sort-seq approach to the development of single fluorescent protein biosensors"

**Supplementary Text. Sequences of the constructs used in this study.**

**PyronicSF-LE-TR**

PdhR 1-188

Linkers

cpGFP

PdhR 199-254

**Amino Acid:**

MGSAYSKIRQPKLSDVIEQQLEFLILEGTLRPGEKLPPERELAKQFDVSRPSLREAIQRLEAKGLLLRRQGGGTFVQSSLWQSFSDPLVELLSDHPESQYDLLETRHALEGIAAYYAALRSTDEDKERIRELHHAIELAQQSGDLDAESNAVLQYQIAVTEAAHNVVLLHLLRCMEPMLAQNVRQNFELLLENVYIKADKQKNGIKANFKIRHNIEDGGVQLAYHYQQNTPIGDGPVLLPDNHYLSVQSKLSKDPNEKRDHMVLLEFVTAAGITLGMDELYKGGTGGSMVSKGEELFTGVVPILVELDGDVNGHKFSVSGEGEGDATYGKLTLKFICTTGKLPVPWPTLVTTLTYGVQCFSRYPDHMKQHDFFKSAMPEGYIQERTIFFKDDGNYKTRAEVKFEGDTLVNRIELKGIDFKEDGNILGHKLEYNTRYSRREMLPLVSSHRTRIFEAIMAGKPEEAREASHRHLAFIEEILLDRSREESRRERSLRRLEQRKNSG*

**DNA:**

ATGGGCAGCGCATATAGCAAAATTCGGCAGCCCAAACTGAGCGATGTGATTGAGCAGCAGCTGGAGTTTCTGATTCTGGAAGGCACCCTGAGGCCTGGAGAGAAACTGCCCCCTGAGCGCGAACTCGCCAAGCAGTTCGACGTGAGTCGACCATCACTGAGGGAGGCTATCCAGAGGCTGGAAGCAAAGGGACTGCTCCTGAGGAGACAGGGAGGAGGGACTTTCGTGCAGAGCTCCCTGTGGCAGAGCTTCAGCGACCCCCTGGTCGAGCTCCTGTCTGACCACCCAGAAAGTCAGTACGATCTCCTGGAGACAAGACATGCTCTGGAAGGCATCGCCGCTTACTATGCAGCCCTGCGGTCCACTGACGAGGATAAGGAACGCATCCGAGAGCTGCACCATGCCATTGAACTCGCTCAGCAGTCAGGAGATCTGGATGCAGAGAGCAACGCCGTGCTGCAGTACCAGATTGCAGTCACCGAGGCTGCACACAATGTGGTCCTCCTGCATCTCCTGAGGTGCATGGAGCCAATGCTGGCCCAGAACGTGAGACAGAATTTTGAGCTCCTGCTCGAAAACGTCTATATCAAGGCCGACAAGCAGAAAAACGGCATTAAGGCTAACTTCAAGATCAGACACAACATCGAGGATGGTGGCGTGCAGCTGGCCTACCATTATCAGCAGAACACACCAATCGGAGATGGACCAGTGCTGCTCCCAGATAATCACTACCTGAGCGTCCAGTCCAAGCTGTCTAAAGACCCTAACGAGAAGCGGGATCATATGGTGCTGCTCGAATTTGTCACAGCCGCTGGGATCACTCTGGGTATGGACGAGCTCTATAAAGGAGGGACCGGTGGCAGTATGGTGTCAAAGGGCGAGGAACTGTTCACAGGAGTGGTCCCCATTCTGGTGGAGCTCGACGGCGATGTCAATGGACACAAATTTTCCGTGTCTGGCGAGGGCGAAGGAGATGCTACCTACGGGAAGCTGACACTCAAATTCATCTGCACCACAGGCAAGCTGCCAGTGCCCTGGCCTACTCTGGTCACTACCCTCACCTACGGGGTGCAGTGTTTCTCCAGATATCCCGACCACATGAAGCAGCATGATTTCTTTAAATCTGCTATGCCTGAGGGGTACATCCAGGAACGGACAATTTTCTTTAAGGACGATGGTAACTACAAAACACGCGCAGAGGTGAAGTTCGAAGGCGACACTCTGGTCAATCGAATCGAGCTGAAGGGAATTGACTTTAAAGAAGATGGGAACATCCTGGGTCACAAGCTGGAGTACAATACTAGGTATTCTCGGCGCGAAATGCTGCCACTCGTGTCTAGTCACAGGACCAGAATCTTTGAGGCAATTATGGCCGGAAAGCCCGAGGAAGCTAGAGAAGCAAGTCACCGGCATCTGGCCTTCATCGAGGAAATTCTGCTCGACCGGAGCCGCGAGGAATCCCGAAGGGAGCGCAGCCTGAGGCGACTCGAACAGCGAAAGAACTCAGGCTAA

**PyronicSF-VE-SS**

PdhR 1-188

Linkers

cpGFP

PdhR 199-254

**Amino Acid:**

MGSAYSKIRQPKLSDVIEQQLEFLILEGTLRPGEKLPPERELAKQFDVSRPSLREAIQRLEAKGLLLRRQGGGTFVQSSLWQSFSDPLVELLSDHPESQYDLLETRHALEGIAAYYAALRSTDEDKERIRELHHAIELAQQSGDLDAESNAVLQYQIAVTEAAHNVVLLHLLRCMEPMLAQNVRQNFELLVENVYIKADKQKNGIKANFKIRHNIEDGGVQLAYHYQQNTPIGDGPVLLPDNHYLSVQSKLSKDPNEKRDHMVLLEFVTAAGITLGMDELYKGGTGGSMVSKGEELFTGVVPILVELDGDVNGHKFSVSGEGEGDATYGKLTLKFICTTGKLPVPWPTLVTTLTYGVQCFSRYPDHMKQHDFFKSAMPEGYIQERTIFFKDDGNYKSSAEVKFEGDTLVNRIELKGIDFKEDGNILGHKLEYNTRYSRREMLPLVSSHRTRIFEAIMAGKPEEAREASHRHLAFIEEILLDRSREESRRERSLRRLEQRKNSG*

**DNA:**

ATGGGCAGCGCATATAGCAAAATTCGGCAGCCCAAACTGAGCGATGTGATTGAGCAGCAGCTGGAGTTTCTGATTCTGGAAGGCACCCTGAGGCCTGGAGAGAAACTGCCCCCTGAGCGCGAACTCGCCAAGCAGTTCGACGTGAGTCGACCATCACTGAGGGAGGCTATCCAGAGGCTGGAAGCAAAGGGACTGCTCCTGAGGAGACAGGGAGGAGGGACTTTCGTGCAGAGCTCCCTGTGGCAGAGCTTCAGCGACCCCCTGGTCGAGCTCCTGTCTGACCACCCAGAAAGTCAGTACGATCTCCTGGAGACAAGACATGCTCTGGAAGGCATCGCCGCTTACTATGCAGCCCTGCGGTCCACTGACGAGGATAAGGAACGCATCCGAGAGCTGCACCATGCCATTGAACTCGCTCAGCAGTCAGGAGATCTGGATGCAGAGAGCAACGCCGTGCTGCAGTACCAGATTGCAGTCACCGAGGCTGCACACAATGTGGTCCTCCTGCATCTCCTGAGGTGCATGGAGCCAATGCTGGCCCAGAACGTGAGACAGAATTTTGAGCTCCTGGTAGAAAACGTCTATATCAAGGCCGACAAGCAGAAAAACGGCATTAAGGCTAACTTCAAGATCAGACACAACATCGAGGATGGTGGCGTGCAGCTGGCCTACCATTATCAGCAGAACACACCAATCGGAGATGGACCAGTGCTGCTCCCAGATAATCACTACCTGAGCGTCCAGTCCAAGCTGTCTAAAGACCCTAACGAGAAGCGGGATCATATGGTGCTGCTCGAATTTGTCACAGCCGCTGGGATCACTCTGGGTATGGACGAGCTCTATAAAGGAGGGACCGGTGGCAGTATGGTGTCAAAGGGCGAGGAACTGTTCACAGGAGTGGTCCCCATTCTGGTGGAGCTCGACGGCGATGTCAATGGACACAAATTTTCCGTGTCTGGCGAGGGCGAAGGAGATGCTACCTACGGGAAGCTGACACTCAAATTCATCTGCACCACAGGCAAGCTGCCAGTGCCCTGGCCTACTCTGGTCACTACCCTCACCTACGGGGTGCAGTGTTTCTCCAGATATCCCGACCACATGAAGCAGCATGATTTCTTTAAATCTGCTATGCCTGAGGGGTACATCCAGGAACGGACAATTTTCTTTAAGGACGATGGTAACTACAAAACACGCGCAGAGGTGAAGTTCGAAGGCGACACTCTGGTCAATCGAATCGAGCTGAAGGGAATTGACTTTAAAGAAGATGGGAACATCCTGGGTCACAAGCTGGAGTACAATAGTAGTTATTCTCGGCGCGAAATGCTGCCACTCGTGTCTAGTCACAGGACCAGAATCTTTGAGGCAATTATGGCCGGAAAGCCCGAGGAAGCTAGAGAAGCAAGTCACCGGCATCTGGCCTTCATCGAGGAAATTCTGCTCGACCGGAGCCGCGAGGAATCCCGAAGGGAGCGCAGCCTGAGGCGACTCGAACAGCGAAAGAACTCAGGCTAA

**PyronicSF-SNAx2/VSTx2**

PdhR 1-188

Linkers

X={A,G,E,L,Q,R,P,V}

Z={A,G,P,R,S,T}

cpGFP

PdhR 199-254

**Amino Acid:**

MGSAYSKIRQPKLSDVIEQQLEFLILEGTLRPGEKLPPERELAKQFDVSRPSLREAIQRLEAKGLLLRRQGGGTFVQSSLWQSFSDPLVELLSDHPESQYDLLETRHALEGIAAYYAALRSTDEDKERIRELHHAIELAQQSGDLDAESNAVLQYQIAVTEAAHNVVLLHLLRCMEPMLAQNVRQNFELLXXNVYIKADKQKNGIKANFKIRHNIEDGGVQLAYHYQQNTPIGDGPVLLPDNHYLSVQSKLSKDPNEKRDHMVLLEFVTAAGITLGMDELYKGGTGGSMVSKGEELFTGVVPILVELDGDVNGHKFSVSGEGEGDATYGKLTLKFICTTGKLPVPWPTLVTTLTYGVQCFSRYPDHMKQHDFFKSAMPEGYIQERTIFFKDDGNYKZZAEVKFEGDTLVNRIELKGIDFKEDGNILGHKLEYNTRYSRREMLPLVSSHRTRIFEAIMAGKPEEAREASHRHLAFIEEILLDRSREESRRERSLRRLEQRKNSG*

**DNA:**

ATGGGCAGCGCATATAGCAAAATTCGGCAGCCCAAACTGAGCGATGTGATTGAGCAGCAGCTGGAGTTTCTGATTCTGGAAGGCACCCTGAGGCCTGGAGAGAAACTGCCCCCTGAGCGCGAACTCGCCAAGCAGTTCGACGTGAGTCGACCATCACTGAGGGAGGCTATCCAGAGGCTGGAAGCAAAGGGACTGCTCCTGAGGAGACAGGGAGGAGGGACTTTCGTGCAGAGCTCCCTGTGGCAGAGCTTCAGCGACCCCCTGGTCGAGCTCCTGTCTGACCACCCAGAAAGTCAGTACGATCTCCTGGAGACAAGACATGCTCTGGAAGGCATCGCCGCTTACTATGCAGCCCTGCGGTCCACTGACGAGGATAAGGAACGCATCCGAGAGCTGCACCATGCCATTGAACTCGCTCAGCAGTCAGGAGATCTGGATGCAGAGAGCAACGCCGTGCTGCAGTACCAGATTGCAGTCACCGAGGCTGCACACAATGTGGTCCTCCTGCATCTCCTGAGGTGCATGGAGCCAATGCTGGCCCAGAACGTGAGACAGAATTTTGAGCTCCTGSNASNAAACGTCTATATCAAGGCCGACAAGCAGAAAAACGGCATTAAGGCTAACTTCAAGATCAGACACAACATCGAGGATGGTGGCGTGCAGCTGGCCTACCATTATCAGCAGAACACACCAATCGGAGATGGACCAGTGCTGCTCCCAGATAATCACTACCTGAGCGTCCAGTCCAAGCTGTCTAAAGACCCTAACGAGAAGCGGGATCATATGGTGCTGCTCGAATTTGTCACAGCCGCTGGGATCACTCTGGGTATGGACGAGCTCTATAAAGGAGGGACCGGTGGCAGTATGGTGTCAAAGGGCGAGGAACTGTTCACAGGAGTGGTCCCCATTCTGGTGGAGCTCGACGGCGATGTCAATGGACACAAATTTTCCGTGTCTGGCGAGGGCGAAGGAGATGCTACCTACGGGAAGCTGACACTCAAATTCATCTGCACCACAGGCAAGCTGCCAGTGCCCTGGCCTACTCTGGTCACTACCCTCACCTACGGGGTGCAGTGTTTCTCCAGATATCCCGACCACATGAAGCAGCATGATTTCTTTAAATCTGCTATGCCTGAGGGGTACATCCAGGAACGGACAATTTTCTTTAAGGACGATGGTAACTACAAAACACGCGCAGAGGTGAAGTTCGAAGGCGACACTCTGGTCAATCGAATCGAGCTGAAGGGAATTGACTTTAAAGAAGATGGGAACATCCTGGGTCACAAGCTGGAGTACAATVSTVSTTATTCTCGGCGCGAAATGCTGCCACTCGTGTCTAGTCACAGGACCAGAATCTTTGAGGCAATTATGGCCGGAAAGCCCGAGGAAGCTAGAGAAGCAAGTCACCGGCATCTGGCCTTCATCGAGGAAATTCTGCTCGACCGGAGCCGCGAGGAATCCCGAAGGGAGCGCAGCCTGAGGCGACTCGAACAGCGAAAGAACTCAGGCTAA

**MBP-162**

MBP 1-162

Linkers

cpGFP

MBP 161-370

**Amino Acid:**

MSKIEEGKLVIWINGDKGYNGLAEVGKKFEKDTGIKVTVEHPDKLEEKFPQVAATGDGPDIIFWAHDRFGGYAQSGLLAEITPDKAFQDKLYPFTWDAVRYNGKLIAYPIAVEALSLIYNKDLLPNPPKTWEEIPALDKELKAKGKSALMFNLQEPYFTWPLIAASYNVFIMADKQKNGIKANFKIRHNIEDGGVQLAYHYQQNTPIGDGPVLLPDNHYLSVQSKLSKDPNEKRDHMVLLEFVTAAGITLGMDELYKGGTGGSMVSKGEELFTGVVPILVELDGDVNGHKFSVSGEGEGDATYGKLTLKFICTTGKLPVPWPTLVTTLTYGVQCFSRYPDHMKQHDFFKSAMPEGYIQERTIFFKDDGNYKTRAEVKFEGDTLVNRIELKGIDFKEDGNILGHKLEYNFNASIAADGGYAFKYENGKYDIKDVGVDNAGAKAGLTFLVDLIKNKHMNADTDYSIAEAAFNKGETAMTINGPWAWSNIDTSKVNYGVTVLPTFKGQPSKPFVGVLSAGINAASPNKELAKEFLENYLLTDEGLEAVNKDKPLGAVALKSYEEELAKDPRIAATMENAQKGEIMPNIPQMSAFWYAVRTAVINAASGRQTVDEALKDAQTRITKSGHHHHHH*

**DNA:**

ATGTCCAAAATCGAAGAAGGTAAACTGGTAATCTGGATTAACGGCGATAAAGGCTATAACGGACTCGCTGAAGTCGGTAAGAAATTCGAGAAAGATACCGGAATTAAAGTCACCGTTGAGCATCCGGATAAACTGGAAGAGAAATTCCCACAGGTTGCGGCAACTGGCGATGGCCCTGACATTATCTTCTGGGCACACGACCGCTTTGGTGGCTACGCTCAATCTGGCCTGTTGGCTGAAATCACCCCGGACAAAGCGTTCCAGGACAAGCTGTATCCGTTTACCTGGGATGCCGTACGTTACAACGGCAAGCTGATTGCTTACCCGATCGCTGTTGAAGCGTTATCGCTGATTTATAACAAAGATCTGCTGCCGAACCCGCCAAAAACCTGGGAAGAGATCCCGGCGCTGGATAAAGAACTGAAAGCGAAAGGTAAGAGCGCGCTGATGTTCAACCTGCAAGAACCGTACTTCACCTGGCCGCTGATTGCTGCATCTTATAACGTCTTTATCATGGCCGACAAGCAGAAGAACGGCATCAAGGCGAACTTCAAGATCCGCCACAACATCGAGGACGGCGGCGTGCAGCTCGCCTATCACTACCAGCAGAACACCCCCATCGGCGACGGCCCCGTGCTGCTGCCCGACAACCACTACCTGAGCGTGCAGTCCAAACTGAGCAAAGACCCCAACGAGAAGCGCGATCACATGGTCCTGCTGGAGTTCGTGACCGCCGCCGGGATCACTCTCGGCATGGACGAGCTGTACAAGGGCGGTACCGGAGGGAGCATGGTGAGCAAGGGCGAGGAGCTGTTCACCGGGGTGGTGCCCATCCTGGTCGAGCTGGACGGCGACGTAAACGGCCACAAGTTCAGCGTGTCCGGCGAGGGCGAGGGCGATGCCACCTACGGCAAGCTGACCCTGAAGTTCATCTGCACCACCGGCAAGCTGCCCGTGCCCTGGCCCACCCTCGTGACCACCCTGACCTACGGCGTGCAGTGCTTCAGCCGCTACCCCGACCACATGAAGCAGCACGACTTCTTCAAGTCCGCCATGCCCGAAGGCTACATTCAGGAGCGCACCATCTTCTTCAAGGACGACGGCAACTATAAGACACGCGCTGAGGTTAAGTTCGAGGGCGACACTCTGGTTAACCGCATCGAGCTGAAGGGCATCGACTTCAAGGAGGACGGCAACATCCTGGGCCATAAGCTTGAATATAACTTCAACGCGTCAATTGCTGCTGACGGGGGTTATGCGTTCAAGTATGAAAACGGCAAGTACGACATTAAAGACGTGGGCGTGGATAACGCTGGCGCGAAAGCGGGTCTGACCTTCCTGGTTGACCTGATTAAAAACAAACACATGAATGCAGACACCGATTACTCCATCGCAGAAGCTGCCTTTAATAAAGGCGAAACAGCGATGACCATCAACGGCCCGTGGGCATGGTCCAACATCGACACCAGCAAAGTGAATTATGGTGTAACGGTACTGCCGACCTTCAAGGGTCAACCATCCAAACCGTTCGTTGGCGTGCTGAGCGCAGGTATTAACGCCGCCAGTCCGAACAAAGAGCTGGCGAAAGAGTTCCTCGAAAACTATCTGCTGACTGATGAAGGTCTGGAAGCGGTTAATAAAGACAAACCGCTGGGTGCCGTAGCGCTGAAGTCTTACGAGGAAGAGTTGGCGAAAGATCCACGTATTGCCGCCACCATGGAAAACGCCCAGAAAGGTGAAATCATGCCGAACATCCCGCAGATGTCCGCTTTCTGGTATGCCGTGCGTACTGCGGTGATCAACGCCGCCAGCGGTCGTCAGACTGTCGATGAAGCCCTGAAAGACGCGCAGACTCGTATCACCAAGAGCGGTCACCATCACCATCACCATTAA

**MBP-164**

MBP 1-164

Linkers

cpGFP

MBP 163-370

**Amino Acid:**

MSKIEEGKLVIWINGDKGYNGLAEVGKKFEKDTGIKVTVEHPDKLEEKFPQVAATGDGPDIIFWAHDRFGGYAQSGLLAEITPDKAFQDKLYPFTWDAVRYNGKLIAYPIAVEALSLIYNKDLLPNPPKTWEEIPALDKELKAKGKSALMFNLQEPYFTWPLIAADASYNVFIMADKQKNGIKANFKIRHNIEDGGVQLAYHYQQNTPIGDGPVLLPDNHYLSVQSKLSKDPNEKRDHMVLLEFVTAAGITLGMDELYKGGTGGSMVSKGEELFTGVVPILVELDGDVNGHKFSVSGEGEGDATYGKLTLKFICTTGKLPVPWPTLVTTLTYGVQCFSRYPDHMKQHDFFKSAMPEGYIQERTIFFKDDGNYKTRAEVKFEGDTLVNRIELKGIDFKEDGNILGHKLEYNFNASADGGYAFKYENGKYDIKDVGVDNAGAKAGLTFLVDLIKNKHMNADTDYSIAEAAFNKGETAMTINGPWAWSNIDTSKVNYGVTVLPTFKGQPSKPFVGVLSAGINAASPNKELAKEFLENYLLTDEGLEAVNKDKPLGAVALKSYEEELAKDPRIAATMENAQKGEIMPNIPQMSAFWYAVRTAVINAASGRQTVDEALKDAQTRITKSGHHHHHH*

**DNA**:

ATGTCCAAAATCGAAGAAGGTAAACTGGTAATCTGGATTAACGGCGATAAAGGCTATAACGGACTCGCTGAAGTCGGTAAGAAATTCGAGAAAGATACCGGAATTAAAGTCACCGTTGAGCATCCGGATAAACTGGAAGAGAAATTCCCACAGGTTGCGGCAACTGGCGATGGCCCTGACATTATCTTCTGGGCACACGACCGCTTTGGTGGCTACGCTCAATCTGGCCTGTTGGCTGAAATCACCCCGGACAAAGCGTTCCAGGACAAGCTGTATCCGTTTACCTGGGATGCCGTACGTTACAACGGCAAGCTGATTGCTTACCCGATCGCTGTTGAAGCGTTATCGCTGATTTATAACAAAGATCTGCTGCCGAACCCGCCAAAAACCTGGGAAGAGATCCCGGCGCTGGATAAAGAACTGAAAGCGAAAGGTAAGAGCGCGCTGATGTTCAACCTGCAAGAACCGTACTTCACCTGGCCGCTGATTGCTGCTGATGCATCTTATAACGTCTTTATCATGGCCGACAAGCAGAAGAACGGCATCAAGGCGAACTTCAAGATCCGCCACAACATCGAGGACGGCGGCGTGCAGCTCGCCTATCACTACCAGCAGAACACCCCCATCGGCGACGGCCCCGTGCTGCTGCCCGACAACCACTACCTGAGCGTGCAGTCCAAACTGAGCAAAGACCCCAACGAGAAGCGCGATCACATGGTCCTGCTGGAGTTCGTGACCGCCGCCGGGATCACTCTCGGCATGGACGAGCTGTACAAGGGCGGTACCGGAGGGAGCATGGTGAGCAAGGGCGAGGAGCTGTTCACCGGGGTGGTGCCCATCCTGGTCGAGCTGGACGGCGACGTAAACGGCCACAAGTTCAGCGTGTCCGGCGAGGGCGAGGGCGATGCCACCTACGGCAAGCTGACCCTGAAGTTCATCTGCACCACCGGCAAGCTGCCCGTGCCCTGGCCCACCCTCGTGACCACCCTGACCTACGGCGTGCAGTGCTTCAGCCGCTACCCCGACCACATGAAGCAGCACGACTTCTTCAAGTCCGCCATGCCCGAAGGCTACATTCAGGAGCGCACCATCTTCTTCAAGGACGACGGCAACTATAAGACACGCGCTGAGGTTAAGTTCGAGGGCGACACTCTGGTTAACCGCATCGAGCTGAAGGGCATCGACTTCAAGGAGGACGGCAACATCCTGGGCCATAAGCTTGAATATAACTTCAACGCGTCAGCTGACGGGGGTTATGCGTTCAAGTATGAAAACGGCAAGTACGACATTAAAGACGTGGGCGTGGATAACGCTGGCGCGAAAGCGGGTCTGACCTTCCTGGTTGACCTGATTAAAAACAAACACATGAATGCAGACACCGATTACTCCATCGCAGAAGCTGCCTTTAATAAAGGCGAAACAGCGATGACCATCAACGGCCCGTGGGCATGGTCCAACATCGACACCAGCAAAGTGAATTATGGTGTAACGGTACTGCCGACCTTCAAGGGTCAACCATCCAAACCGTTCGTTGGCGTGCTGAGCGCAGGTATTAACGCCGCCAGTCCGAACAAAGAGCTGGCGAAAGAGTTCCTCGAAAACTATCTGCTGACTGATGAAGGTCTGGAAGCGGTTAATAAAGACAAACCGCTGGGTGCCGTAGCGCTGAAGTCTTACGAGGAAGAGTTGGCGAAAGATCCACGTATTGCCGCCACCATGGAAAACGCCCAGAAAGGTGAAATCATGCCGAACATCCCGCAGATGTCCGCTTTCTGGTATGCCGTGCGTACTGCGGTGATCAACGCCGCCAGCGGTCGTCAGACTGTCGATGAAGCCCTGAAAGACGCGCAGACTCGTATCACCAAGAGCGGTCACCATCACCATCACCATTAA

**EMMA-attB-Dest**

**Source:**

HC_Kan_attB-p1

YCe2982 HC_Kan_conn_B-E

YCe3756 HC_Kan_conn_E-F

YCe1895 HC_Kan_Kozak-ATG_p6

YCe1695 HC_Kan_RFP-p7 **(BsmbI sites)**

YCe1921 HC_Kan_conn_H-K

YCe1889 HC_Kan_SV40-polyA_p11

YCe2731 HC_Kan_conn_L-W

YCe2721 HC_Kan_conn_W-Z

YCe3736_HC_Amp_ccdB_receiver vector

**Sequence:**

TAGGAAGGCTTGTCGACGACGGCGGTCTCCGTCGTCAGGATCATCGATGGACTAACTACGGTGAAAGAGACTTAGAAGGAGATTGAGAGATCCGTTAGCCTTTAGTGAACCGTCAGAATTAATTCAGATCGATCTACCAGAACCGTCAGATCCGCTAGAGATTACGCCAACCGCCACCATGGGCAGCA**GAGACG**GAATTCGCGGCCGCTTCTAGAGCAATACGCAAACCGCCTCTCCCCGCGCGTTGGCCGATTCATTAATGCAGCTGGCACGACAGGTTTCCCGACTGGAAAGCGGGCAGTGAGCGCAACGCAATTAATGTGAGTTAGCTCACTCATTAGGCACCCCAGGCTTTACACTTTATGCTTCCGGCTCGTATGTTGTGTGGAATTGTGAGCGGATAACAATTTCACACATACTAGAGAAAGAGGAGAAATACTAGATGGCTTCCTCCGAAGACGTTATCAAAGAGTTCATGCGTTTCAAAGTTCGTATGGAAGGTTCCGTTAACGGTCACGAGTTCGAAATCGAAGGTGAAGGTGAAGGTCGTCCGTACGAAGGTACCCAGACCGCTAAACTGAAAGTTACCAAAGGTGGTCCGCTGCCGTTCGCTTGGGACATCCTGTCCCCGCAGTTCCAGTACGGTTCCAAAGCTTACGTTAAACACCCGGCTGACATCCCGGACTACCTGAAACTGTCCTTCCCGGAAGGTTTCAAATGGGAACGTGTTATGAACTTCGAAGACGGTGGTGTTGTTACCGTTACCCAGGACTCCTCCCTGCAAGACGGTGAGTTCATCTACAAAGTTAAACTGCGTGGTACCAACTTCCCGTCCGACGGTCCGGTTATGCAGAAAAAAACCATGGGTTGGGAAGCTTCCACCGAACGTATGTACCCGGAAGACGGTGCTCTGAAAGGTGAAATCAAAATGCGTCTGAAACTGAAAGACGGTGGTCACTACGACGCTGAAGTTAAAACCACCTACATGGCTAAAAAACCGGTTCAGCTGCCGGGTGCTTACAAAACCGACATCAAACTGGACATCACCTCCCACAACGAAGACTACACCATCGTTGAACAGTACGAACGTGCTGAAGGTCGTCACTCCACCGGTGCTTAATAACGCTGATAGTGCTAGTGTAGATCGCTACTAGAGCCAGGCATCAAATAAAACGAAAGGCTCAGTCGAAAGACTGGGCCTTTCGTTTTATCTGTTGTTTGTCGGTGAACGCTCTCTACTAGAGTCACACTGGCTCACCTTCGGGTGGGCCTTTCTGCGTTTATATACTAGTAGCGGCCGCTGCAG**CGTCTC**TAGGCTAATAACAGCTTCCGGACTCTAGAACATCCCTACAGGTGATATCCTCGGGTAACTTGTTTATTGCAGCTTATAATGGTTACAAATAAAGCAATAGCATCACAAATTTCACAAATAAAGCATTTTTTTCACTGCATTCTAGTTGTGGTTTGTCCAAACTCATCAATGTATCTTATCATGTCTGTCGTCTATGATGAGGATGTTGGTGGAGAGCATGTGGAGGAAGTGGATAGGGAAGGTTGTAGAGTAGATCCGGTTGAAGTGATGAGGATAGGAGGAGGCGAACACATATTGATCCTCGTCACAACCACACCAGCACACCTCAAATCCCACACCACTCCCACAATTACCATTCACTCAACAAACTCACACATCCCACGATAACGAATTCAAGCTTGATATCATTCAGGACGAGCCTCAGACTCCAGCGTAACTGGACTGCAATCAACTCACTGGCTCACCTTCACGGGTGGGCCTTTCTTCGGTAGAAAATCAAAGGATCTTCTTGAGATCCTTTTTTTCTGCGCGTAATCTGCTGCTTGCAAACAAAAAAACCACCGCTACCAGCGGTGGTTTGTTTGCCGGATCAAGAGCTACCAACTCTTTTTCCGAGGTAACTGGCTTCAGCAGAGCGCAGATACCAAATACTGTTCTTCTAGTGTAGCCGTAGTTAGGCCACCACTTCAAGAACTCTGTAGCACCGCCTACATACCTCGCTCTGCTAATCCTGTTACCAGTGGCTGCTGCCAGTGGCGATAAGTCGTGTCTTACCGGGTTGGACTCAAGACGATAGTTACCGGATAAGGCGCAGCGGTCGGGCTGAACGGGGGGTTCGTGCACACAGCCCAGCTTGGAGCGAACGACCTACACCGAACTGAGATACCTACAGCGTGAGCTATGAGAAAGCGCCACGCTTCCCGAAGGGAGAAAGGCGGACAGGTATCCGGTAAGCGGCAGGGTCGGAACAGGAGAGCGCACGAGGGAGCTTCCAGGGGGAAACGCCTGGTATCTTTATAGTCCTGTCGGGTTTCGCCACCTCTGACTTGAGCATCGATTTTTGTGATGCTCGTCAGGGGGGCGGAGCCTATGGAAAAACGCCAGCAACGCAGAAAGGCCCACCCGAAGGTGAGCCAGGTGATTACATTTGGGCCCTCATTACCAATGCTTAATCAGTGAGGCACCTATCTCAGCGATCTGTCTATTTCGTTCATCCATAGTTGCCTGACTCCCCGTCGTGTAGATAACTACGATGCGGGAGGGCTTACCATCTGGCCCCAGTGCTGCAATGATACCGCGAGAACCACGCTCACCGGCTCCAGATTTATCAGCAATAAACCAGCCAGCCGGGAGGGCCGAGCGCAGAAGTGATCCTGCAACTTTATCCGCCTCCATCCAGTCTATTAATTGTTGCCGGGAAGCTAGAGTAAGTAGTTCGCCAGTTAATAGTTTGCGCAACGTTGTTGCCATTGCTACAGGCATCGTGGTGTCACGCTCGTCGTTTGGTATGGCTTCATTCAGCTCCGGTTCCCAACGATCAAGGCGAGTTACATGATCCCCCATGTTGTGCAAAAAAGCGGTTAGCTCCTTCGGTCCTCCGATCGTTGCCAGAAGTAAGTTGGCCGCAGTGTTATCACTCATGGTTATGGCAGCACTGCATAATTCTCTTACTGTCATGCCATCCGTGAGATGCTTTTCTGTGACTGGTGAGTACTCAACCAAGTCATTCTGAGAATAGTGTATGCGGCGACCGAGTTGCTCTTGCCCGGCGTCAATACGGGATAATACCGCGCCACATAGCAGAACTTTAAAAGTGCTCATCATTGGAAAACGTTCTTCGGGGCGTAAACTCTCAAGGATCTTACCGCTGTTGAGATCCAGTTCGATGTAACCCACTCGTGCACCCAACTGATCTTCAGCATCTTTTACTTTCACCAGCGTTTCTGGGTGAGCAAAAACAGGAAGGCAAAATGCCGCAAAAAAGGGAATAAGGGCGACACGGAAATGTTGAATACTCATTTTAGCTTCCTTAGCTCCTGAAAATCTCGATAACTCAAAAAATACGCCCGGTAGTGATCTTATTTCATTATGGTGAAAGTTGGAACCTCTTACGTGCCGATCAAGTCAAAAGCCTCCGGTCGGAGGCTTTTGACTTTCTGCTATGGAGGTCAGGTATGATTTAAATGGTCAGTATTGAGCGATATCTAGAGAATTCGTCA
